# GatorAnchor: Cross-Modal Uncertainty–Aware Learning of Reliable Cell Neighborhoods in Single-Cell Multi-Omics

**DOI:** 10.64898/2026.11.14.766684

**Authors:** Zhenhao Zhang, Youhong Tang, Jiang Bian, Xiangtao Li, Yuxi Liu

## Abstract

Cross-modal uncertainty challenges single-cell multi-omics integration because molecular modalities differ in sparsity, technical noise, and context-dependent biological informativeness, producing discordant cell–cell relationships. Treating conflicting evidence as equally reliable can propagate uncertain relationships and obscure biologically meaningful cellular heterogeneity. GatorAnchor is a cross-modal uncertainty-aware framework that learns modality-specific probabilistic representations, quantifies disagreement across modalities and neighborhood evidence sources, and refines cell–cell relationships according to their reliability. Across ten paired transcriptome–protein and transcriptome–chromatin datasets, GatorAnchor achieves the highest mean performance across four evaluation metrics among nine representative integration methods, demonstrating improved resolution of cellular heterogeneity. Candidate relations with greater composite unreliability show lower consistency with reference cell-type structure, whereas filtering highly unreliable relations increases same-type edge purity across all datasets. Modality conflict is associated with transcript–protein measurement discordance, while transcriptome–chromatin analyses reveal lineage-dependent contributions of molecular and decoder-derived evidence. A three-modality transcriptome– chromatin–protein analysis characterizes cluster of differentiation 4 (CD4)-related communities with distinct molecular profiles and consistent within-donor contrasts across 12 donors. Under incomplete and mosaic measurements, GatorAnchor preserves informative cell-population structure, while cross-modal correspondence increasingly depends on paired-cell anchors. These findings support uncertainty-aware neighborhood refinement for reliable and interpretable characterization of cellular heterogeneity.

## Introduction

Recent advances in single-cell multi-omics profiling have enabled multiple molecular layers to be measured simultaneously within the same cell, providing a detailed view of cellular identity, state and regulation^1–3^.

Gene expression, cell-surface protein abundance and chromatin accessibility capture distinct dimensions of transcriptional activity, cellular phenotype and regulatory state, enabling cellular heterogeneity to be characterized more comprehensively than with any individual assay^4–6^. Integrating these complementary molecular layers can further connect cellular phenotypes with their underlying regulatory programs, providing a more comprehensive basis for identifying disease-associated cell states and interpreting molecular mechanisms^7–14^. However, these modalities differ in feature dimensionality, sparsity, signal distribution and susceptibility to technical noise^15^. Their relative informativeness can also vary across cellular contexts, with different modalities contributing unequally to the resolution of particular cell populations^16,17^. Accordingly, the same cells can show different local neighborhood structures across modality-specific representations^16^. This variability creates cross-modal uncertainty in multimodal integration: cellular organization must be learned without assuming uniform reliability across molecular modalities or the cell–cell relationships inferred from them.

Many existing single-cell multi-omics integration methods address cross-modal heterogeneity by learning shared or aligned low-dimensional representations. Factorization-based approaches such as MOFA+ model variation that is shared or specific across modalities and sample groups^18^, whereas scMoMaT uses matrix tri-factorization for mosaic integration, learning cell representations together with modality-specific feature representations^19^. Deep generative approaches provide probabilistic alternatives: scMM combines modality-specific variational posteriors through a mixture-of-experts formulation^20^, while MIDAS uses product-of-experts inference together with self-supervised modality alignment and disentanglement of biological state from technical variation^16^. More recent methods, including scMultiNet^21^ and scECDA^22^, incorporate attention-based fusion, contrastive learning and denoising to improve cross-modal representation learning.

These approaches have advanced the integration of heterogeneous multimodal measurements by modeling shared structure, modality-specific variation, technical noise and adaptive feature weighting. However, their objectives primarily focus on representation learning and cross-modal alignment; modality-specific uncertainty and cross-modal disagreement are not typically translated into relation-specific assessments that determine which cell–cell connections should be trusted for information propagation.

Graph and relational approaches explicitly exploit relational structure during multimodal integration^15,23,24^. scDMSC learns self-expression coefficients in a shared latent space and derives a symmetric cell–cell similarity matrix for spectral clustering^23^. DeepMAPS constructs a heterogeneous cell–gene graph and applies a heterogeneous graph transformer to jointly learn cell and gene representations^24^, whereas scMAGCA constructs a KNN cell graph from concatenated, preprocessed multimodal features and uses graph convolution to learn topology-preserving cell representations^15^. These approaches demonstrate the value of relational information for multimodal representation learning and clustering. Yet the benefit of graph-based propagation depends on whether the edges used for message passing are sufficiently reliable^25–27^. A cell–cell relation supported by one modality but contradicted by another may propagate information between cells whose local association is uncertain. Such unreliable propagation may blur meaningful differences among cellular populations and states, thereby limiting the characterization of cellular heterogeneity. Thus, an unresolved question is not only how to construct or weight a multimodal graph, but also how to determine which cell–cell relations should be trusted for information propagation.

Addressing this question requires distinguishing modality-specific uncertainty from cross-modal disagreement, two related but conceptually different sources of unreliability. A modality may yield a low-uncertainty representation while another modality supports a substantially different representation of the same cell; confidence within individual modalities therefore does not imply cross-modal agreement. The same principle extends to the neighborhood level, where multiple evidence sources—RNA-derived similarity, ADT- or ATAC-derived similarity, and learned edge-decoder support—may provide conflicting evidence for the same cell–cell relation. Rather than treating such disagreement only as a discrepancy to be minimized during alignment, it can provide information about the reliability of the relation itself. This motivates explicitly modeling both source-specific uncertainty and cross-modal or cross-source disagreement when determining which neighborhood relations should be trusted, allowing multimodal integration to preserve reliable local structure without obscuring meaningful cellular heterogeneity.

Here, we present GatorAnchor, a cross-modal uncertainty-aware framework for learning reliable cell neighborhoods in paired single-cell multi-omics data. At the representation level, GatorAnchor learns modality-specific Gaussian representations from graph-augmented features and combines them through precision-weighted product-of-experts fusion. Because PoE variance reflects the uncertainty of the individual experts but does not directly capture disagreement between their latent means, GatorAnchor explicitly incorporates cross-modal mean disagreement into the fused latent dispersion. At the neighborhood level, after an initial warm-up stage, candidate cell–cell relations are assembled from the observation graph, modality-specific neighborhoods and the learned fused representation. For each candidate relation, RNA similarity, second-modality similarity and shared edge-decoder support are integrated through uncertainty-aware source weighting, while disagreement among these evidence sources contributes to a composite edge-unreliability score. Relations with greater unreliability are attenuated or removed to construct a refined propagation graph. This graph is refined once after warm-up and subsequently held fixed during residual graph adaptation and joint fine-tuning.

Across ten paired RNA+ADT and RNA+ATAC datasets, GatorAnchor achieved the highest mean AMI, NMI, ARI and ACC compared with nine representative single-cell multi-omics integration methods. Edge-reliability analyses showed that candidate relations assigned greater composite unreliability were less consistent with reference cell-type structure, while filtering highly unreliable relations increased same-type edge purity across all ten datasets, with dataset-dependent effects on graph connectivity. In RNA+ADT analyses, modality conflict was associated with RNA–protein measurement discordance, and high-confidence edges showed greater molecular similarity and cross-modal neighborhood consistency. A paired RNA+ATAC PBMC analysis, GatorAnchor recovered concordant lineage-associated transcriptional and chromatin-accessibility programs, while the relative contributions of RNA, ATAC and decoder-derived evidence varied across cellular lineages. We further applied GatorAnchor to jointly measured RNA, ATAC and ADT data, where it characterized CD4-related communities with molecular profiles and consistent within-donor molecular contrasts. Under incomplete and mosaic measurements, GatorAnchor preserved informative cell-population structure, while cross-modal correspondence became increasingly dependent on paired-cell anchors. These results support explicit modeling of neighborhood reliability as a strategy for improving multimodal integration while providing interpretable diagnostics of cross-modal agreement and graph evidence.

## Results

### Overview of the GatorAnchor framework

Paired single-cell multi-omics assays measure multiple molecular layers in the same cells. However, differences in sparsity, noise, and signal quality can make molecular evidence unequally reliable across modalities and produce discordant cell–cell neighborhood structures. GatorAnchor addresses this cross-modal uncertainty by modeling both modality-specific uncertainty and cross-modal disagreement, rather than assuming that molecular evidence or the resulting cell–cell relationships are uniformly reliable. The framework first constructs a measurement-derived observation graph and learns modality-specific probabilistic representations from graph-augmented features. These representations are combined through precision-weighted product-of-experts fusion, with cross-modal latent disagreement incorporated into the fused uncertainty. After an initial warm-up stage, candidate cell–cell relations are assembled from the observation graph, the fused latent representation, and modality-specific neighborhoods. Each candidate relation is then evaluated using RNA similarity, second-modality similarity, and support from a shared edge decoder. These evidence sources are integrated using uncertainty-derived weights, while disagreement among them contributes to a composite edge-level unreliability score. Relations with greater unreliability are attenuated or removed to construct a refined propagation graph. Graph refinement is performed once after warm-up, and the resulting graph is held fixed during residual graph convolutional network adaptation and subsequent joint fine-tuning. An overview of the GatorAnchor framework is shown in **Figure 1**.

**Figure 1.**
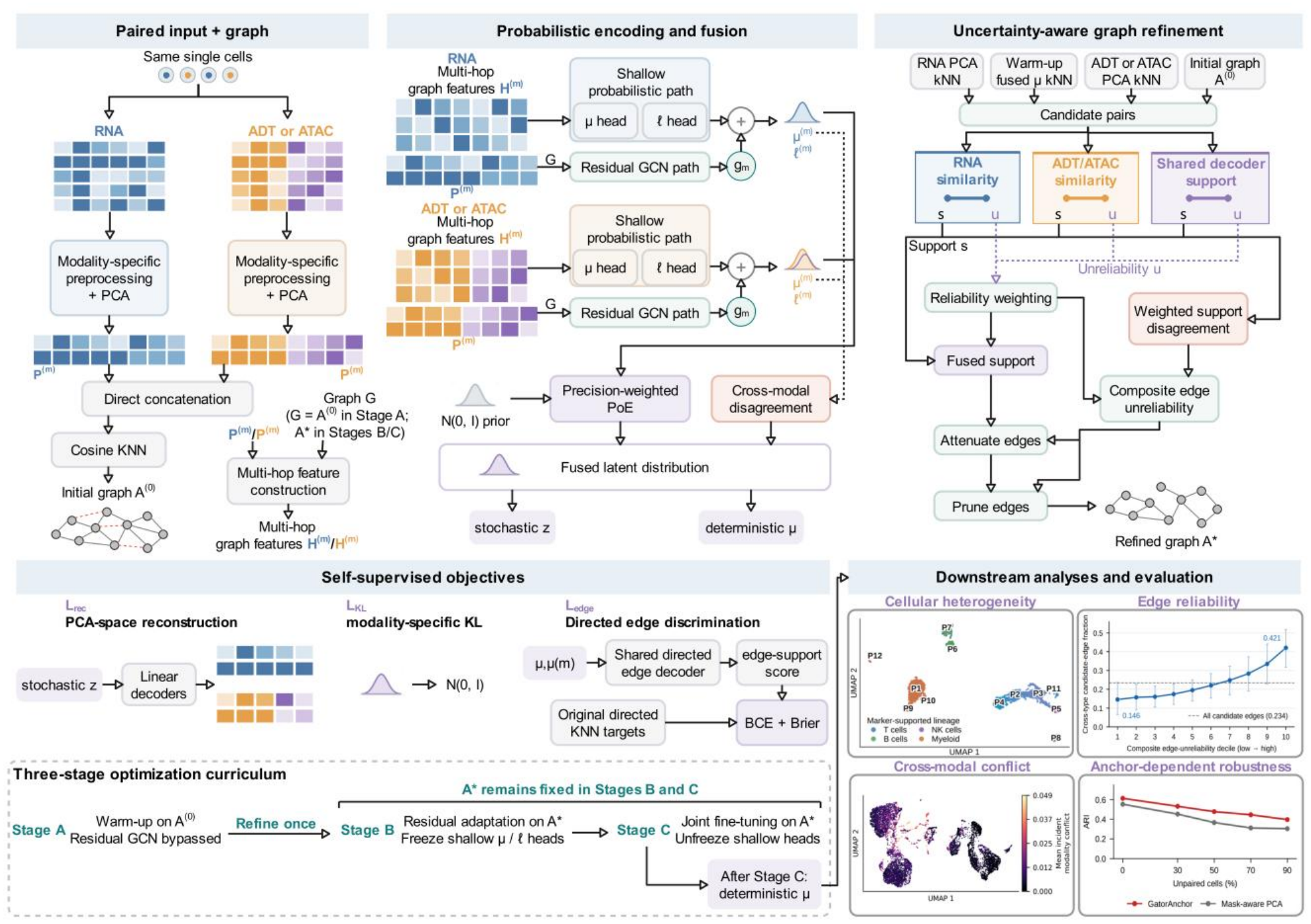
Overview of the GatorAnchor framework.

### Benchmark datasets, data preprocessing, and baseline methods

We evaluated our proposed GatorAnchor on ten publicly available paired single-cell multi-omics datasets, comprising five paired RNA and antibody-derived tag (RNA+ADT) datasets and five RNA and chromatin accessibility (RNA+ATAC) datasets. RNA+ADT datasets included 10Xmalt, GSE128639^34^, GSE194122 s3d6, 10X 1k PBMC, and GSE158013^35^. RNA+ATAC datasets included GSE194122 s4d8, GSE201402^36^, PBMC unsorted 3k, GSE156478 Stim^37^, and Brain SNARE^38^. These datasets were generated using diverse single-cell multi-omics technologies, including CITE-seq, 10x Feature Barcoding, TEA-seq, 10x Multiome, DOGMA-seq, and SNARE-seq. For the TEA-seq and DOGMA-seq datasets, we analyzed the RNA+ADT and RNA+ATAC modality pairs, respectively.

Specifically, dataset sizes ranged from 713 to 30,672 cells, with 5 to 36 annotated cell types per dataset. The input matrices contained 6,275–36,601 RNA features, 17–134 ADT features, and 58,376–230,281 ATAC peaks. For datasets containing more than two molecular modalities, we selected two modalities to maintain a consistent paired multi-omics setting: RNA+ADT for GSE158013 and RNA+ATAC for GSE156478 Stim. Cell-type annotations provided with the datasets were used as reference labels for evaluating clustering performance. Detailed characteristics of the ten benchmark datasets are summarized in **Table 1**.

**Table 1.** Statistics of the ten paired single-cell multi-omics datasets used.

| Dataset | No. of cells | Omics1 |  | Omics2 |  | Reference classes | Technology | Reference |
|---|---|---|---|---|---|---|---|---|
|  |  | Modality | No. of genes | Modality | No. of features |  |  |  |
| 10Xmalt | 8,412 | RNA | 33,538 | ADT | 17 | 11 | 10x Feature Barcoding | -- |
| GSE128639 | 30,672 | RNA | 17,009 | ADT | 25 | 27 | CITE-seq | [34] |
| GSE194122 s3d6 | 11,035 | RNA | 13,953 | ADT | 134 | 36 | 10x Feature Barcoding | -- |
| 10X 1k PBMC | 713 | RNA | 33,538 | ADT | 17 | 5 | 10x Feature Barcoding | -- |
| GSE158013 | 25,517 | RNA | 36,601 | ADT | 46 | 12 | TEA-seq | [35] |
| GSE194122 s4d8 | 9,876 | RNA | 13,431 | ATAC | 113,600 | 19 | 10x Multiome | -- |
| GSE201402 | 9,383 | RNA | 6,275 | ATAC | 58,376 | 16 | 10x Multiome | [36] |
| PBMC unsorted 3k | 2,413 | RNA | 36,601 | ATAC | 78,218 | 12 | 10x Multiome | -- |
| GSE156478 Stim | 5,915 | RNA | 36,495 | ATAC | 66,476 | 14 | DOGMA-seq | [37] |
| Brain SNARE | 8,055 | RNA | 33,160 | ATAC | 230,281 | 13 | SNARE-seq | [38] |

For GatorAnchor, RNA and the second modality were processed independently for each dataset using Scanpy^39^. Measurements were normalized to a total count of 10^4^ per cell and log-transformed using log(1 + *x*). For RNA and ATAC, a predefined number of highly variable genes or peaks was retained when the number of available features exceeded the target, whereas all available ADT features were retained. The retained features were standardized and separately projected using modality-specific PCA, generating the score matrices P^(*r*)^ and P^(2)^ defined in the Methods section.

We compared GatorAnchor against nine representative single-cell multi-omics integration methods spanning probabilistic factor analysis and matrix factorization^18,19^, deep generative modeling^16,20^, graph-based representation learning^15,24^, adversarial learning^15,21^, attention-based multimodal fusion^21,22^, and deep multi-view subspace learning^23^.

**MOFA+**^18^ is a Bayesian group factor analysis framework that combines automatic relevance determination, sparsity-inducing priors, and stochastic variational inference to disentangle variation that is shared or specific across data modalities and sample groups.

**scMM**^20^ is a mixture-of-experts multimodal variational autoencoder that combines modality-specific variational posteriors into a joint latent representation and enables bidirectional cross-modal generation through modality-specific decoders.

**DeepMAPS**^24^ represents integrated multi-omics data as a heterogeneous graph of cell and gene nodes and applies a multi-head heterogeneous graph transformer to jointly learn cell and gene embeddings for cell clustering and cell-type-specific biological network inference.

**scMoMaT**^19^ performs mosaic integration using matrix tri-factorization to integrate an arbitrary number of single-cell data matrices, learning cell and modality-specific feature representations while identifying cluster-specific biomarkers across modalities.

**MIDAS**^16^ is a VAE-based deep generative framework for mosaic integration that combines self-supervised modality alignment with information-theoretic disentanglement of biological state and technical noise, enabling dimensionality reduction, imputation, and batch correction.

**scMAGCA**^15^ constructs a KNN cell graph and combines graph convolution, adversarial learning, MSE and zero-inflated negative binomial reconstruction, and KL-divergence-based clustering refinement to learn topology-preserving cell embeddings in an end-to-end framework.

**scMultiNet**^21^ is a multi-task deep adversarial neural network built on an autoencoder architecture that combines bidirectional cross-modal prediction, contrastive learning, multi-head self-attention, and indicator-guided reconstruction for joint denoising, multimodal integration, and cross-modal prediction.

**scECDA**^22^ uses omics-specific autoencoders, enhanced contrastive learning with latent-space perturbation-based augmentation, and differential-attention feature fusion to align and denoise cross-omics representations and directly produce integrated latent features and cell-clustering assignments.

**scDMSC**^23^ combines weighted multimodal reconstruction with self-expression-based deep multi-view subspace learning to obtain shared latent features and a similarity matrix derived from self-expression coefficients for subsequent spectral clustering.

### GatorAnchor accurately resolves cellular heterogeneity across paired single-cell multi-omics datasets

To evaluate the ability of GatorAnchor to resolve cellular heterogeneity, we benchmarked it against nine representative single-cell multi-omics integration methods across ten paired single-cell multi-omics datasets. Clustering performance was evaluated using AMI, NMI, ARI, and ACC. As shown in **Figure 2**, GatorAnchor achieved the highest overall mean performance across the ten datasets for all four metrics. Notable improvements were observed on several RNA+ATAC datasets. On Brain SNARE, GatorAnchor achieved AMI, NMI, ARI, and ACC scores of 0.7947, 0.7956, 0.7764, and 0.8752, respectively, outperforming scMAGCA, the strongest baseline on this dataset across all four metrics, by 0.1126, 0.1121, 0.2586, and 0.2814. On PBMC unsorted 3k, GatorAnchor achieved an ARI of 0.8577 and an ACC of 0.8850, compared with 0.6280 and 0.7287 for scDMSC, the best-performing baseline for these two metrics. Consistent gains were also observed on the RNA+ADT datasets. On GSE128639, GatorAnchor obtained AMI, NMI, ARI, and ACC scores of 0.8781, 0.8785, 0.9010, and 0.8529, respectively, compared with 0.8308, 0.8316, 0.8434, and 0.8018 for scMultiNet, which was the strongest baseline across all four metrics on this dataset. Collectively, these results demonstrate that GatorAnchor consistently improves the identification of annotated cell populations across diverse paired single-cell multi-omics datasets.

**Figure 2.**
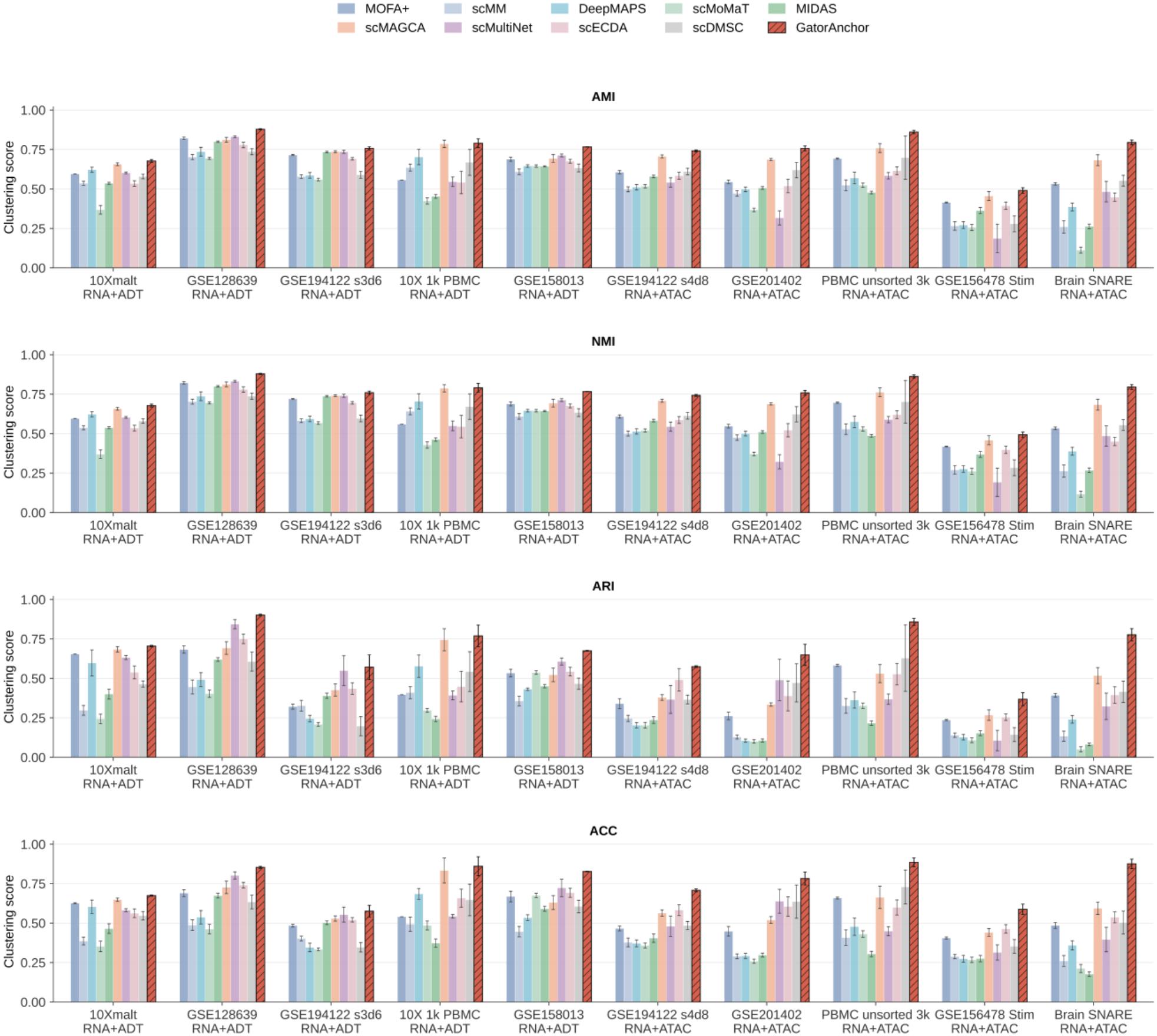
Benchmarking GatorAnchor for identification of cellular populations across paired single-cell multi-omics datasets. Performance was evaluated using AMI, NMI, ARI, and ACC. Bars show mean scores across repeated runs, with error bars indicating standard deviations.

### Composite edge unreliability provides a decomposable graph-reliability diagnostic

We evaluated whether composite edge unreliability provides an informative diagnostic of candidate cell–cell relationships rather than serving only as an internal graph-weighting quantity. Specifically, we examined whether higher unreliability ranks candidate neighborhoods that are less consistent with reference cell-type structure, whether filtering these edges improves local graph purity while characterizing its structural consequences, and whether edge unreliability can be decomposed into distinct modality- and reconstruction-related disagreement patterns. We further assessed the computational cost of the uncertainty diagnostic and the sensitivity of downstream clustering to a strict observation-edge holdout protocol (**Figure 3**). Candidate edges included observation-graph relations together with local relationships proposed from the fused and assay-specific representations. Reference cell-type annotations were used only for post-hoc evaluation.

**Figure 3.**
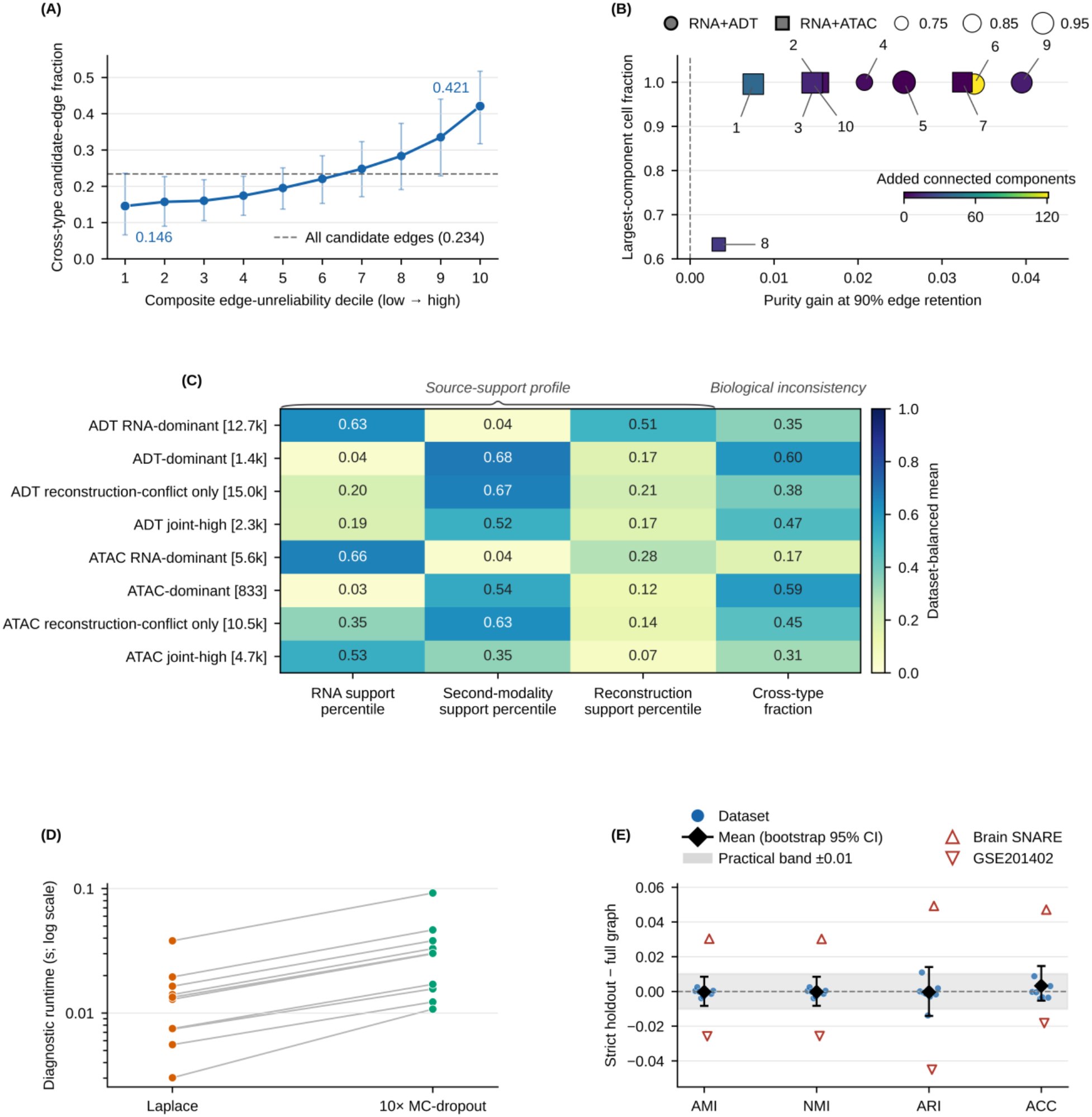
Decomposable edge-reliability diagnostics and graph refinement in GatorAnchor. (A) Cross-type candidate-edge fraction across within-run deciles of composite edge unreliability. The dashed line denotes the dataset-balanced fraction across all candidate edges. (B) Purity and graph-structure changes after retaining the 90% least-unreliable candidate edges. The axes show same-type purity gain and largest-component cell fraction; marker shape denotes assay type, marker size denotes rare-type degree retention, and color denotes added connected components. (C) Source-support profiles of RNA-dominant, second-modality-dominant, reconstruction-conflict-only, and joint-high conflict edges in RNA+ADT and RNA+ATAC datasets. The first three columns show source-wise support percentiles and the final column shows cross-type edge fraction; brackets indicate typical candidate-edge counts. (D) Paired diagnostic runtimes of the last-layer Laplace approximation and ten-pass Monte Carlo (MC) dropout across datasets. (E) Changes in AMI, NMI, ARI, and ACC under strict edge holdout relative to full-observation-graph training. Points denote datasets, diamonds and error bars denote dataset-balanced means and bootstrap 95% confidence intervals, and shading indicates the prespecified ±0.01 practical band. GSE201402 and Brain SNARE are highlighted because each has at least one metric with |Δ| > 0.02.

We assessed whether composite unreliability meaningfully ranks candidate edges according to biological neighborhood inconsistency (**Figure 3A**). Candidate edges were stratified into within-run unreliability deciles, from the most reliable to the least reliable. The dataset-balanced fraction of edges connecting different reference cell types increased from 0.146 in the lowest-unreliability decile to 0.421 in the highest, corresponding to a D10−D1 difference of 0.275 (bootstrap 95% CI, 0.140–0.403; BH-adjusted *q* = 0.027). The association was positive in 9 of 10 datasets, with a median within-dataset Spearman *ρ* of 0.970 (BH-adjusted *q* = 0.027). These results show that candidate edges assigned higher composite unreliability are enriched for neighborhoods that are less consistent with reference cell-type structure. Cross-type status was treated as a proxy for biological neighborhood inconsistency rather than as a definitive edge-error label, because biologically meaningful transitions or closely related cell states can also generate cross-type connections.

We assessed whether this ranking can guide graph filtering without achieving higher purity simply through excessive edge removal (**Figure 3B**). We retained the 90% least-unreliable candidate edges and evaluated both same-type edge purity and graph preservation. Same-type purity increased in all ten datasets, with a dataset-balanced mean gain of 0.0208 (95% CI, 0.0139–0.0276). The structural consequences, however, were dataset dependent. PBMC unsorted 3k retained only 0.633 of cells in its largest connected component, indicating substantial fragmentation, whereas GSE128639 retained a largest-component fraction of 0.996 but generated approximately 121 additional connected components. Thus, preservation of the dominant component can coexist with fragmentation of smaller cell subsets, and higher edge purity does not necessarily imply a structurally preferable graph. The composite score should therefore not be interpreted as a universally superior edge-ranking score; its additional value lies in the ability to decompose unreliability into interpretable sources of evidence disagreement.

We therefore examined which forms of evidence disagreement characterize unreliable candidate edges (**Figure 3C**). Direction- and assay-stratified profiles separated disagreement between RNA and the second modality from disagreement between observation-derived and decoder-derived edge support. Second-modality-dominant conflicts showed the highest cross-type fractions in both RNA+ADT and RNA+ATAC datasets (0.60 and 0.59, respectively), indicating that candidate edges with stronger ADT or ATAC support but weaker RNA support were particularly enriched across reference cell-type boundaries. Reconstruction-conflict-only profiles represented a different pattern, with relatively stronger second-modality support and weaker decoder-derived support: the corresponding support percentiles were 0.67 versus 0.21 in RNA+ADT and 0.63 versus 0.14 in RNA+ATAC. Joint-high profiles showed simultaneous elevation of modality and reconstruction conflict but differed between assay types. These patterns indicate that the composite score distinguishes qualitatively different forms of source disagreement.

Because this diagnostic is evaluated over a large candidate-edge set, we assessed its computational cost relative to repeated MC-dropout estimation (**Figure 3D**). The last-layer Laplace approximation was faster than ten-pass Monte Carlo (MC) dropout in all ten datasets. Median dataset-level runtimes were 0.0132 and 0.0301 s, respectively, and MC dropout required a median 2.32 times as long as Laplace (paired Wilcoxon test on dataset-level log-ratios, BH-adjusted *q* = 0.009). The absolute difference between the two medians was approximately 0.017 s.

We examined whether reuse of observation edges during model fitting could influence the reliability analysis. To assess sensitivity to this source of information reuse, we compared a connectivity-preserving strict edge-holdout protocol with full-observation-graph training (**Figure 3E**). The dataset-balanced mean strict-holdout-minus-full-graph differences were −0.00025 for AMI, −0.00024 for NMI, −0.00036 for ARI, and 0.00324 for ACC, indicating no systematic average degradation under the fixed holdout protocol. Dataset-level effects were heterogeneous: GSE201402 decreased across all four metrics, including an ARI change of −0.0450, whereas Brain SNARE showed increases, including an ARI change of 0.0492. Thus, observation-edge holdout did not introduce a consistent average performance loss, but individual datasets differed in their sensitivity to graph-edge perturbation.

Overall, these results support composite edge unreliability as a decomposable graph-reliability diagnostic that ranks biologically inconsistent candidate neighborhoods and separates distinct sources of edge disagreement. Its value lies in combining edge-level ranking with source-specific interpretation rather than serving as a calibrated edge-error probability.

### GatorAnchor modality conflict tracks cross-modal measurement discordance and neighborhood reliability

We assessed whether GatorAnchor modality conflict provides an interpretable measure of cross-modal reliability rather than serving only as an internal graph diagnostic. We therefore examined whether higher conflict corresponded to greater RNA–ADT discordance, whether this relationship persisted after excluding the evaluated features from model fitting, whether it depended on correct RNA–ADT pairing, and whether low-conflict, low-uncertainty edges formed more coherent multimodal neighborhoods. In the 10Xmalt dataset, the integrated embedding organized the benchmark-provided numeric reference classes P1–P11 into several compact regions, whereas modality conflict remained heterogeneous within multiple classes and formed localized hotspots (**Figure 4A,B**). Thus, the spatial distribution of conflict was not determined by reference-class identity alone.

**Figure 4.**
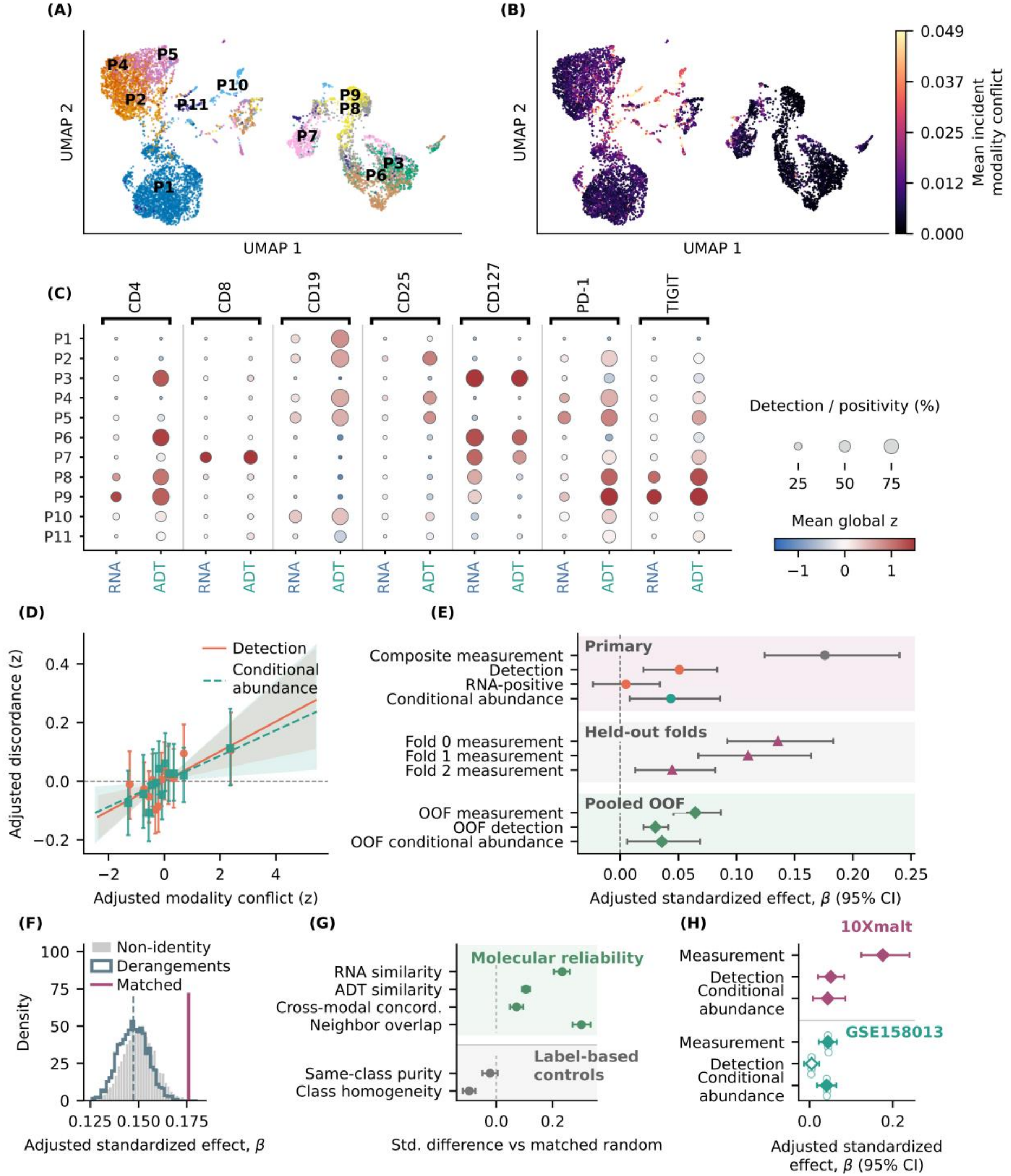
GatorAnchor modality conflict tracks cross-modal measurement discordance and neighborhood reliability. (A) Integrated embedding of 10Xmalt cells colored by benchmark-provided reference classes P1–P11. (B) Cell-level RNA–ADT modality conflict mapped onto the same embedding. (C) RNA and ADT profiles of seven matched marker pairs. Dot size indicates detection or positivity rate, and color indicates mean standardized abundance. (D) Covariate-adjusted associations of modality conflict with detection and conditional abundance discordance. (E) Effect estimates from the complete-feature analysis, feature-held-out folds, and pooled out-of-feature (OOF) analysis. (F) Matched-pair specificity analysis comparing the correct RNA–ADT pairing with non-identity mappings and complete derangements. (G) Molecular consistency and reference-label controls for high-confidence edges relative to matched random edges. Horizontal bars indicate 95% matched-randomization intervals. (H) Directional cross-dataset validation in 10Xmalt and GSE158013. Horizontal bars indicate 95% block-bootstrap confidence intervals, and small open circles denote GSE158013 training-seed sensitivity estimates. ADT, antibody-derived tag; OOF, out of feature.

We tested whether modality conflict tracked disagreement between paired RNA and protein measurements. Seven RNA–ADT pairs—CD4, CD8, CD19, CD25, CD127, PD-1, and TIGIT—were selected from a prespecified ten-pair candidate panel (**Figure 4C**). Detection discordance quantified disagreement between RNA-detection and ADT-positivity states, whereas conditional abundance discordance quantified standardized RNA–ADT differences among observations positive in both modalities. After adjustment for RNA and ADT library size, detected genes, IgG background, candidate degree, and numeric reference class, higher modality conflict was associated with both detection discordance (*β* = 0.0509, 95% block-bootstrap CI 0.0200–0.0832) and conditional abundance discordance (*β* = 0.0435, 95% CI 0.0083–0.0858; both BH-adjusted *q* < 0.05; **Figure 4D**). The latter effect was less precise because fewer cells satisfied the joint-detection criterion.

**Figure 4E** further assessed the robustness of this association to direct feature reuse. The complete-feature composite measurement outcome showed the largest effect (*β* = 0.1759, 95% CI 0.1240–0.2400), whereas the RNA-positive sensitivity outcome was close to zero (*β* = 0.0050, 95% CI −0.0233–0.0341). We then retrained GatorAnchor in three prespecified feature-held-out folds after excluding the corresponding RNA and ADT features before preprocessing and representation learning. The held-out composite-measurement effects remained positive but heterogeneous across folds (0.1355, 0.1099, and 0.0447). In the pooled out-of-feature (OOF) analysis, modality conflict remained positively associated with composite measurement (*β* = 0.0646, 95% CI 0.0457–0.0865), detection (*β* = 0.0303, 95% CI 0.0201–0.0413), and conditional abundance discordance (*β* = 0.0358, 95% CI 0.0060–0.0686). Thus, the association persisted after feature exclusion but was attenuated and marker dependent.

We tested whether correct RNA–ADT correspondence contributed information beyond this broad reliability signal. Across all 5,039 non-identity mappings, the mean adjusted effect was 0.1504, and across the 1,854 complete derangements retaining no correct pair, the mean was 0.1473 (**Figure 4F**). The correctly matched panel yielded an effect of 0.1759, exceeding the derangement mean by 0.0286 (*P* = 0.000539). Because mismatched panels still retained substantial positive associations, the dominant signal reflects broad cross-modal measurement reliability, with a smaller component attributable to correct RNA–ADT pairing.

We assessed whether low-conflict, low-uncertainty edges represented more coherent multimodal neighborhoods. Relative to stringently matched random candidate edges, the 3,295 high-confidence edges showed greater RNA similarity (0.234), ADT similarity (0.105), cross-modal concordance (0.072), and RNA–ADT neighborhood overlap (0.302; **Figure 4G**). Same-class purity did not differ clearly from matched controls, and local reference-class homogeneity was lower. These results indicate that high-confidence edges capture molecular and cross-modal neighborhood consistency rather than simply favoring cells with the same reference label.

We evaluated the same relationships in the GSE158013 RNA+ADT dataset (**Figure 4H**). Modality conflict remained positively associated with composite measurement discordance (*β* = 0.0440, 95% block-bootstrap CI 0.0221–0.0653) and conditional abundance discordance (*β* = 0.0408, 95% CI 0.0175–0.0640), whereas the detection estimate was small and inconclusive (*β* = 0.0049, 95% CI −0.0135–0.0232). Measurement and conditional-abundance effects were directionally stable across three training seeds. Because the eligible marker panels differed between datasets, this analysis provides directional cross-dataset construct validation rather than exact same-panel replication. Overall, these results support GatorAnchor modality conflict as an interpretable diagnostic of cross-modal measurement and molecular-neighborhood reliability.

### GatorAnchor recovers biologically coherent PBMC organization and reveals interpretable cross-modal graph evidence

We examined whether the cellular organization learned by GatorAnchor was supported by concordant signals across molecular modalities and whether its model-defined graph-evidence quantities were interpretable in paired RNA–ATAC data. We focused on the PBMC unsorted 3k dataset, which contains 2,413 cells jointly profiled for 36,601 RNA features and 78,218 chromatin-accessibility peaks and is accompanied by 12 numeric reference classes. Using the fixed GatorAnchor representation, we analyzed the resulting cell groups, modality-specific marker profiles, and post-hoc graph diagnostics. Reference labels were used only after representation learning to specify the target number of clusters, evaluate the partition, and select candidates.

The frozen integrated representation organized the cells into 12 visually separated predicted clusters (**Figure 5A**). To compare this partition with the available reference classes, we constructed a row-normalized reference-by-prediction contingency matrix and used Hungarian matching to align and reorder predicted-cluster labels for visualization (**Figure 5B**). Reference identifiers 0–11 are displayed as R1–R12, whereas predicted clusters are shown as the Hungarian-aligned labels P1–P12. Most reference classes were dominated by a single predicted cluster, producing a pronounced near-diagonal correspondence pattern. The clearest local split occurred for R5, with 80.0% of its cells assigned to P5 and 18.6% assigned to P3. Across the complete partition, GatorAnchor achieved an ARI of 0.892, NMI of 0.884, AMI of 0.883, and Hungarian-matched ACC of 0.938. Thus, the frozen representation closely recovered the supplied reference partition.

**Figure 5.**
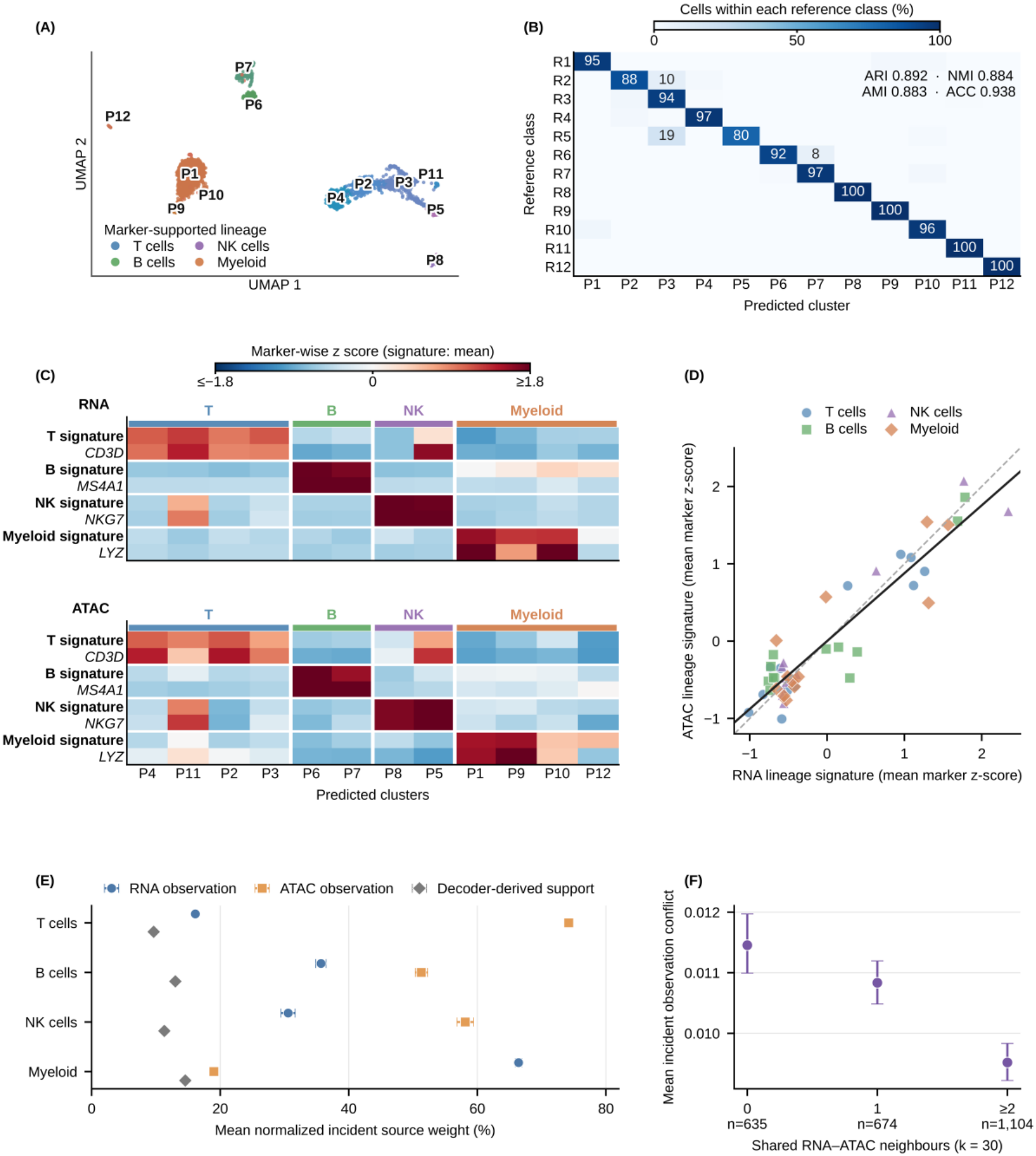
Reference-class recovery, lineage-associated RNA–ATAC programs, and graph-evidence diagnostics of GatorAnchor on PBMC unsorted 3k. The analysis was performed using the frozen GatorAnchor representation and its 12-cluster partition of 2,413 cells. R1–R12 denote one-based reference-class labels, and P1–P12 denote Hungarian-aligned predicted-cluster display labels. (A) UMAP of the integrated representation, colored by post-hoc marker-supported broad lineages; labels indicate predicted clusters. (B) Row-normalized correspondence between reference classes and predicted clusters. Colors indicate the percentage of cells within each reference class assigned to each predicted cluster; ARI, NMI, AMI, and ACC summarize partition agreement. (C) RNA-abundance and ATAC-derived gene-level chromatin-accessibility profiles for T-cell, B-cell, NK-cell, and myeloid signatures across predicted clusters. Representative constituent markers (*CD3D, MS4A1, NKG7*, and *LYZ*) are shown alongside the corresponding signatures. (D) Concordance between RNA and ATAC lineage signatures across 48 lineage-signature–cluster observations (Spearman *ρ* = 0.708, **95% CI 0.626–0.787; permutation** *P* = 0.0002). The dashed line denotes identity and the solid line shows the overall trend. (E) Mean normalized incident weights of RNA observation, ATAC observation, and decoder-derived edge support across marker-supported lineages. Points denote lineage means and horizontal bars indicate 95% bootstrap intervals. (F) Mean incident RNA–ATAC observation conflict stratified by the number of shared neighbors between RNA-PCA and ATAC-PCA 30-nearest-neighbor sets. Points denote group means and vertical bars indicate 95% bootstrap intervals.

We assessed whether the predicted clusters were supported by concordant immune-lineage programs across RNA abundance and chromatin accessibility (**Figure 5C**). Eighteen predefined markers were evaluated using normalized RNA abundance and gene-level chromatin-accessibility scores derived from ATAC peaks using the MAESTRO-Enhanced scheme. Within each modality, each marker was standardized across the 12 predicted clusters. T-cell, B-cell, NK-cell, and myeloid signatures were calculated as the mean marker-wise *z*-score of their constituent markers, and the corresponding RNA and ATAC signatures were averaged to obtain a joint score used to group and order the displayed clusters. The heatmaps additionally show one representative constituent marker for each lineage signature.

The resulting profiles supported a broad lineage organization in both modalities. The strongest joint T-cell signatures occurred in P4, P11, P2, and P3; B-cell signatures in P6 and P7; NK-cell signatures in P8 and P5; and myeloid signatures in P1, P9, P10, and P12. Some clusters retained mixed or weak secondary programs. In particular, P11 showed a strong T-cell signature together with NK/cytotoxic-associated signal, whereas the small P12 cluster (*n* = 18) showed only weak joint myeloid support. These profiles indicate concordant broad lineage-associated structure across the RNA and ATAC modalities.

Across the four lineage signatures and 12 predicted clusters, RNA and ATAC signature values were positively associated (**Figure 5D**). The 48 lineage-signature–cluster observations yielded a Spearman correlation of *ρ* = 0.708 (95% lineage-block bootstrap interval, 0.626–0.787; two-sided within-lineage cluster-label permutation *P* = 0.0002). Leave-one-lineage-out correlations remained positive (0.648–0.735), indicating that the association was not driven by a single lineage. These results indicate that the broad lineage structure recovered by GatorAnchor was consistently reflected across transcriptional and chromatin-accessibility programs.

We examined whether the composition of graph evidence varied across marker-supported lineages (**Figure 5E**). ATAC observation received the largest mean weight in the T-cell (74.22%), B-cell (51.29%), and NK-cell (58.12%) groups, whereas RNA observation predominated in the myeloid group (66.42%). Decoder-derived edge support contributed a smaller but consistently retained component across all four lineages. These patterns indicate that the relative composition of graph evidence varies across marker-supported cellular contexts rather than following a uniform modality mixture.

We assessed whether RNA–ATAC observation conflict varied with agreement between modality-specific neighborhoods (**Figure 5F**). Separate 30-nearest-neighbor sets were constructed in RNA-PCA and ATAC-PCA space, and cells with greater neighborhood overlap showed progressively lower observation conflict. Using the unbinned shared-neighbor count, neighborhood agreement was negatively associated with observation conflict (Spearman *ρ* = −0.230), with mean conflict approximately 17% lower in cells sharing at least two neighbors than in cells with no shared neighbors. This result supports the observation-conflict diagnostic as an indicator of local disagreement between RNA- and ATAC-derived neighborhoods.

Overall, **Figure 5** shows that the frozen GatorAnchor representation closely recapitulated the PBMC reference partition and was accompanied by concordant RNA and ATAC lineage-associated signals. The graph diagnostics further revealed lineage-dependent evidence weighting, while cells with greater RNA– ATAC neighborhood overlap showed lower observation-conflict scores. These results complement the clustering benchmarks by demonstrating both the biological coherence of the learned representation and the interpretability of its graph-evidence diagnostics.

### GatorAnchor shows anchor-dependent robustness under incomplete and mosaic modalities

We evaluated GatorAnchor under three missingness settings to assess clustering, dependence on paired cells, and recovery of masked-population structure (**Figure 6A**). One-sided missingness retained RNA for all cells while withholding the second modality from a subset. The reciprocal mosaic setting comprised RNA-only, second-modality-only, and fully paired cells. Type-block missingness withheld the second modality from an entire prespecified cell population. Each cell retained at least one observed modality throughout these experiments. The unpaired-cell fraction denotes the proportion of cells with exactly one unobserved modality; thus, 90% unpaired cells corresponded to approximately 10% paired cells. The same fixed masks were shared across compared methods, and within each incomplete-data run, only measurements retained by that run’s mask were used. Reciprocal mosaic, paired-anchor and type-block analyses used 10Xmalt and GSE158013 (RNA+ADT), and PBMC unsorted 3k and GSE201402 (RNA+ATAC).

**Figure 6.**
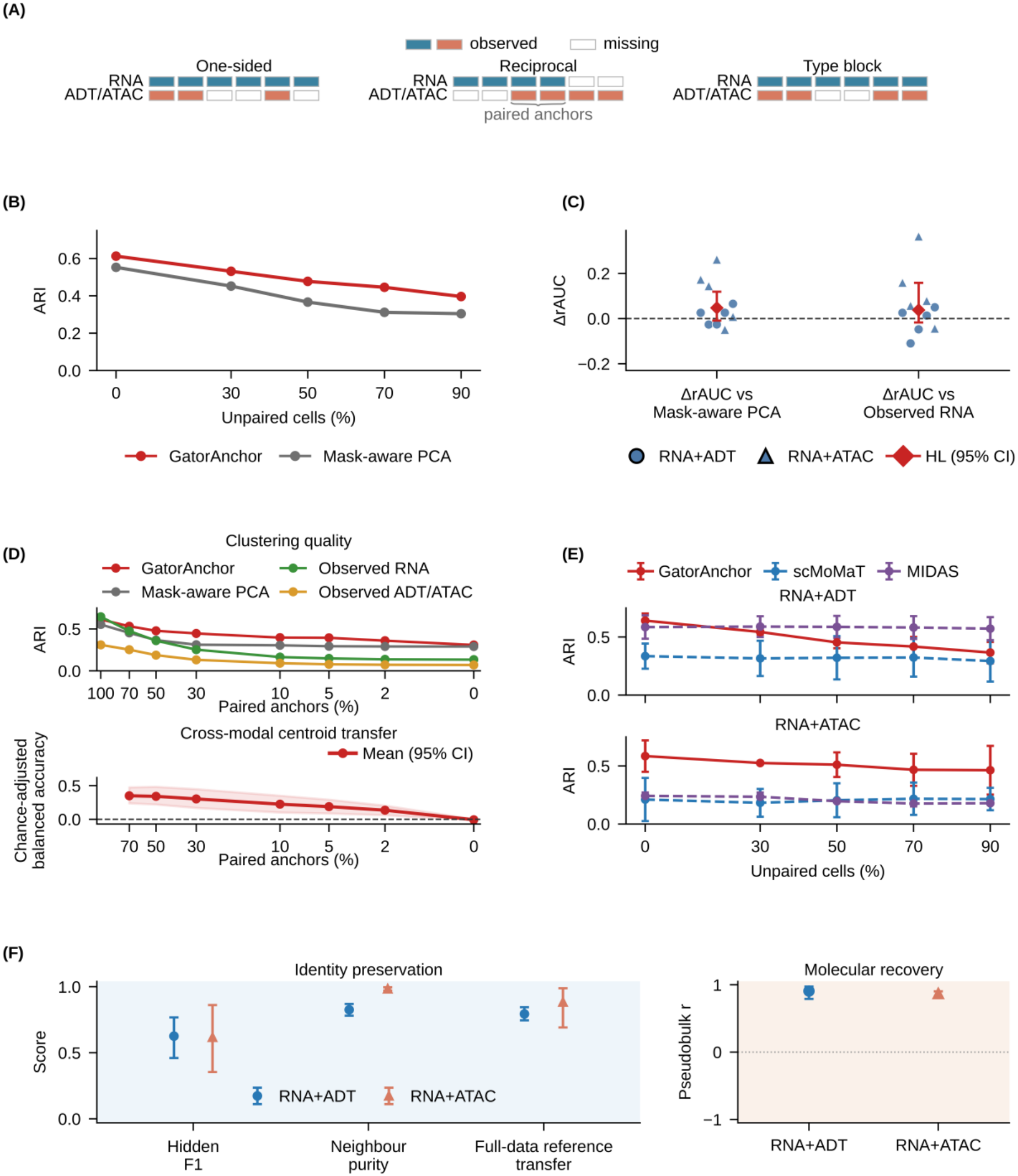
Anchor-dependent robustness of GatorAnchor under incomplete and mosaic modalities. Panels B, D, E and F used 10Xmalt and GSE158013 (RNA+ADT), and PBMC unsorted 3k and GSE201402 (RNA+ATAC). Panel C used all ten datasets listed in Table 1. (A) One-sided, reciprocal mosaic, and type-block missingness designs; filled and open boxes indicate observed and unobserved modalities, respectively. (B) Mean ARI for GatorAnchor and Mask-aware PCA across four datasets under reciprocal mosaic missingness; the x-axis denotes the percentage of unpaired cells. (C) Dataset-level differences in normalized ARI robustness area under the curve (ΔrAUC) relative to Mask-aware PCA and observed RNA across ten one-sided datasets. Circles and triangles denote RNA+ADT and RNA+ATAC datasets, and diamonds show Hodges–Lehmann estimates with 95% confidence intervals. (D) Clustering performance and cross-modal centroid transfer across decreasing paired-anchor fractions. (E) Mean ARI for GatorAnchor, scMoMaT, and MIDAS under reciprocal mosaic missingness across two RNA+ADT and two RNA+ATAC datasets. (F) Masked-population F1, same-type neighborhood purity, full-data reference transfer accuracy, and selected-feature pseudobulk recovery across 20 type-block experiments. Transfer used matched fully observed references with self-matches excluded.

Under reciprocal mosaic missingness, GatorAnchor achieved higher mean ARI than Mask-aware PCA at every tested paired-anchor fraction across four datasets (**Figure 6B**). As paired cells decreased from 100% to 10%, mean ARI declined from 0.613 to 0.396 for GatorAnchor and from 0.553 to 0.304 for Mask-aware PCA. In the one-sided analysis across ten datasets, normalized ARI robustness area under the curve (rAUC) favored GatorAnchor in seven datasets against each baseline (**Figure 6C**). Datasets served as the independent units for these paired comparisons. Hodges–Lehmann paired-difference estimates were 0.047 relative to Mask-aware PCA and 0.038 relative to the observed-RNA baseline. However, the confidence intervals included zero, and neither comparison was significant after Benjamini–Hochberg correction (both *q* = 0.52).

We further reduced the paired-anchor fraction to zero to examine clustering and cross-modal centroid transfer (**Figure 6D**). GatorAnchor retained a mean ARI of 0.309 without paired cells. Chance-adjusted balanced accuracy for centroid transfer declined from 0.349 at 70% paired cells to −0.007 at zero paired cells. Thus, centroid transfer approached chance level in the tested zero-anchor setting despite residual clustering structure. This result distinguishes preserved clustering structure from successful cross-modal correspondence under this evaluation protocol.

We compared GatorAnchor with scMoMaT [19] and MIDAS [16] under reciprocal mosaic missingness across two RNA+ADT and two RNA+ATAC datasets (**Figure 6E**). At 90% unpaired cells, MIDAS achieved the highest mean ARI across the two RNA+ADT datasets, whereas GatorAnchor achieved the highest mean ARI across the two RNA+ATAC datasets. These rankings differed between the assay groups in this four-dataset benchmark.

Across 20 prespecified type-block experiments, mean F1 scores for the modality-masked population were 0.626 for RNA+ADT and 0.617 for RNA+ATAC (**Figure 6F**). We also evaluated same-type neighborhood purity and label-transfer accuracy for the modality-masked cells using fully observed runs of the same cells as references. Each query’s own reference counterpart was excluded before nearest-neighbor voting. Mean transfer accuracies were 0.794 for RNA+ADT and 0.883 for RNA+ATAC. This analysis assessed post-hoc agreement with a full-data reference from the same cells rather than generalization to independent query samples. Selected-feature pseudobulk recovery yielded mean Pearson correlations of 0.907 for RNA+ADT and 0.869 for RNA+ATAC, reflecting population-level recovery of the selected features.

Overall, these results showed retained clustering structure under modality missingness and reduced centroid-transfer accuracy as paired anchors were removed.

### Ablation analysis reveals contributions of GatorAnchor components

To assess the contribution of the major components of GatorAnchor, we constructed six ablation variants and evaluated them across the ten paired single-cell multi-omics datasets (**Figure 7**). Within the multi-source edge fusion module, **F1** replaced uncertainty-aware fusion with a naive equal-weight graph union, **F2** removed confidence gating, and **F3** excluded cross-source conflict from the fused uncertainty. Within the deep residual branch, **D4** removed the residual-branch adaptation stage and directly jointly optimized the shallow and deep branches after graph refinement, whereas **D5** replaced graph convolution with a parameter-matched residual MLP, removing neighborhood propagation within the deep branch. Finally, **S6** removed the Brier-score term from the edge-prediction objective, leaving binary cross-entropy as the sole edge loss. For each variant, performance changes were defined as Δ = metric_variant_ − metric_GatorAnchor_, such that negative values indicate reduced performance relative to the full model. Each model was evaluated across ten seeds per dataset, and dataset-specific changes were calculated from seed-averaged scores.

**Figure 7.**
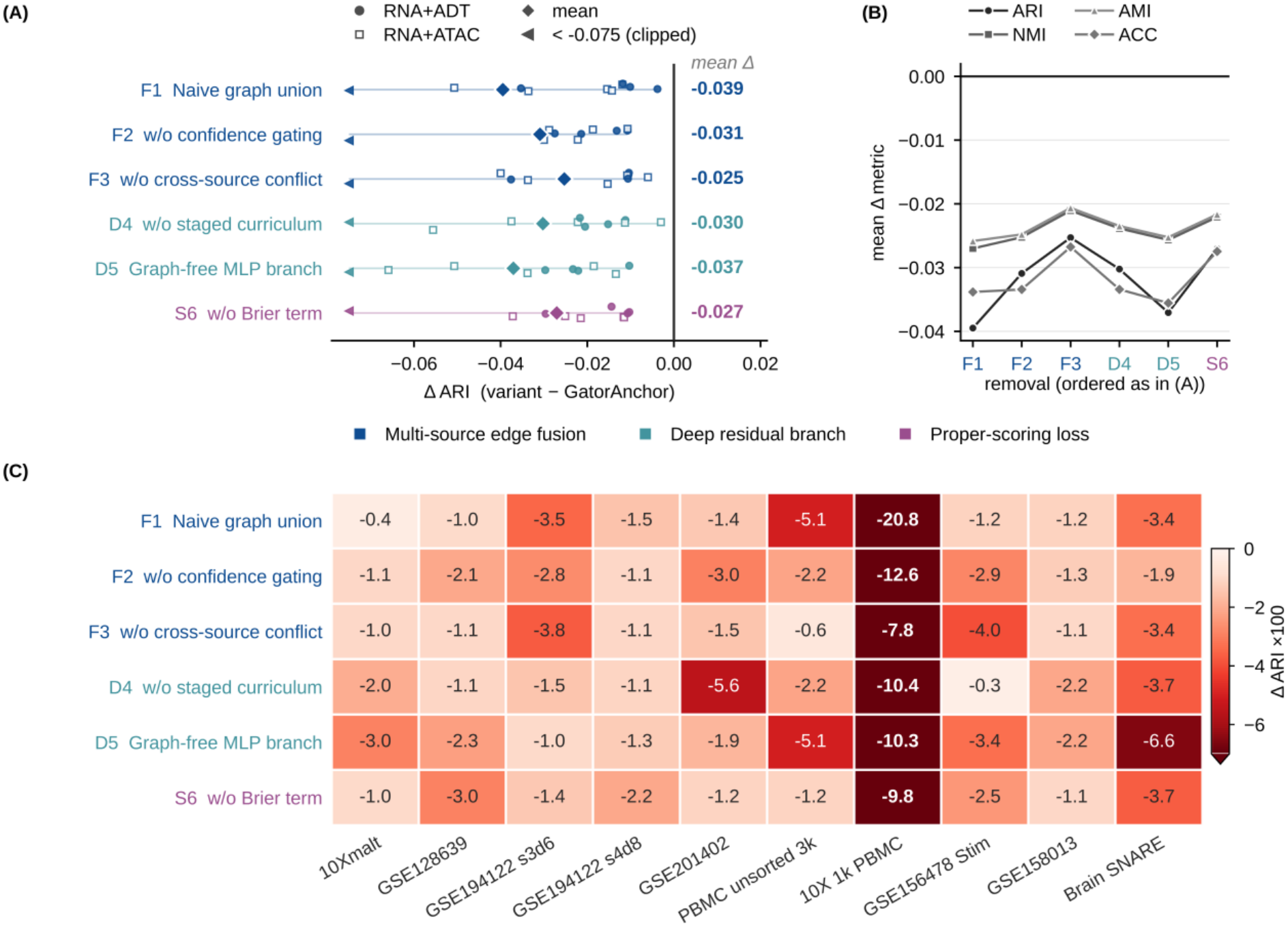
Ablation analysis of GatorAnchor across ten paired single-cell multi-omics datasets. (A) Dataset-specific changes in ARI for six ablation variants relative to the full GatorAnchor model. Filled circles and open squares denote RNA+ADT and RNA+ATAC datasets, respectively, whereas diamonds indicate the mean ΔARI across the ten datasets for each ablation variant. Left-pointing triangles indicate values below the clipping threshold (ΔARI < −0.075). (B) Mean changes in ARI, NMI, AMI, and ACC across the ten datasets. (C) Heatmap of dataset-specific ΔARI values, displayed as ΔARI× 100 for visualization. Negative values indicate reduced performance relative to the full model.

As shown in **Figure 7A**, every ablation reduced ARI across all ten datasets, yielding negative ΔARI values for all 60 variant–dataset comparisons. The largest average decrease was observed for **F1**, with an average ΔARI of −0.0395. A comparable reduction was observed for **D5** (ΔARI=-0.0371), indicating that replacing the graph-based deep branch with a parameter-matched MLP consistently impaired performance. Removing confidence gating (**F2**) and the residual-branch adaptation stage (**D4**) resulted in average ΔARI values of −0.0309 and −0.0302, respectively. Consistent reductions were also observed after removing the Brier term (**S6**, −0.0270) and cross-source conflict modeling (**F3**, −0.0253). This trend was consistent across all four clustering metrics. **Figure 7B** shows that the mean changes in ARI, NMI, AMI, and ACC across the ten datasets were negative for every ablation variant. Panels A and B report metric differences on their original scales, whereas **Figure 7C** displays dataset-specific ΔARI values multiplied by 100 for visualization. The heatmap shows that all 60 variant–dataset combinations had negative mean ΔARI values, confirming that no ablation improved mean ARI on any of the ten datasets.

Two ablations further clarify the contributions of the corresponding model components. In **F1**, the multi-source candidate information was retained while uncertainty-aware evidence fusion was replaced by simple graph aggregation. The resulting performance decrease indicates that the benefit of multi-source integration depends not only on combining multiple graphs, but also on appropriately weighting and filtering heterogeneous edge evidence according to uncertainty. The performance reduction for **D5** supports a contribution of neighborhood propagation within the residual branch beyond parameter count alone. The consistent performance decrease observed for **S6** further indicates that the Brier component of the edge-prediction objective also contributes to downstream clustering performance.

### GatorAnchor shows robust performance across the tested hyperparameter ranges

To assess hyperparameter sensitivity, we evaluated GatorAnchor across ten paired single-cell multi-omics datasets (**Figure 8**). We examined graph neighborhood size (*k*_graph_), propagation order (hops), KL regularization weight (*β*), PoE conflict coefficient (*λ*_PoE_), and uncertainty-gating temperature (*T*). Here, *k*_graph_ and *λ*_PoE_ correspond to *k*_0_ and *λ*_*D*_, respectively, in Methods. For each hyperparameter, one parameter was varied over five values while the remaining parameters were held fixed. Clustering performance was evaluated using AMI, NMI, ARI, and ACC, with each point denoting the mean across five seeds.

**Figure 8.**
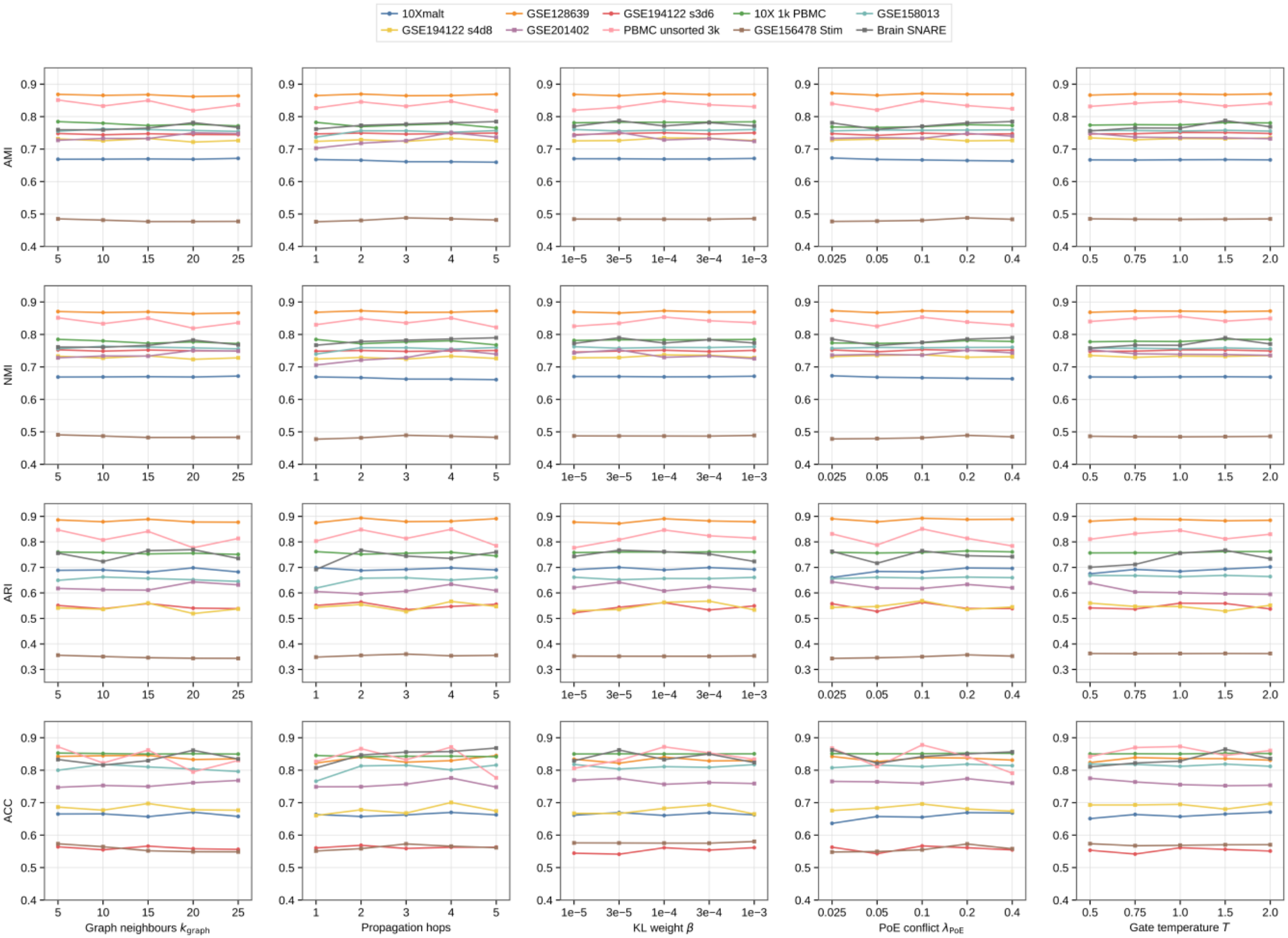
Hyperparameter sensitivity of GatorAnchor across ten paired single-cell multi-omics datasets. Rows show AMI, NMI, ARI, and ACC, and columns correspond to *k*_graph_, propagation order, *β, λ*_PoE_, and *T*. Each curve represents one dataset, with points showing the mean across five seeds.

Propagation order produced the largest average within-dataset variation across the four metrics. The preferred propagation order varied across datasets and evaluation metrics, and increasing propagation order did not consistently improve performance. The remaining hyperparameters produced smaller average within-dataset changes over the tested ranges. Responses to *k*_graph_, *β*, and *λ*_PoE_ were non-monotonic, with dataset-specific optima. Among the tested hyperparameters, *T* produced the smallest average within-dataset variation across all four metrics. Dataset-averaged performance was similar at *T* = 1.0 and *T* = 1.5.

### GatorAnchor characterizes CD4-related communities across three molecular modalities

To examine the applicability of GatorAnchor beyond dual-modality integration, we analyzed jointly measured RNA, ATAC, and ADT profiles from 46,532 CD4-related cells across 12 donors in the ALTRA TEA-seq dataset^40^. The cohort included all cells assigned to the publisher’s CD4 Naive, CD4 Memory, Treg, and CD4 CTL reference categories. Model-based clustering of the final tri-modal representation using mclust yielded four predicted communities, P1–P4, containing 24,380, 19,185, 2,592, and 375 cells, respectively (**Figure 9A**). The predicted communities corresponded predominantly to Naive, Memory, Treg, and CD4 CTL reference cells, respectively (**Figure 9B**). Overall agreement was ARI = 0.8247, NMI = 0.7360, AMI = 0.7359, and ACC = 0.9440. In the row-normalized reference-by-community matrix, 96.31% of Naive cells mapped to P1, 93.43% of Memory cells to P2, 84.09% of Treg cells to P3, and 98.03% of CD4 CTL cells to P4. These percentages quantify reference-class recall rather than predicted-community purity. Treg showed the least complete correspondence, with the remaining 15.91% distributed mainly between P1 and P2.

**Figure 9.**
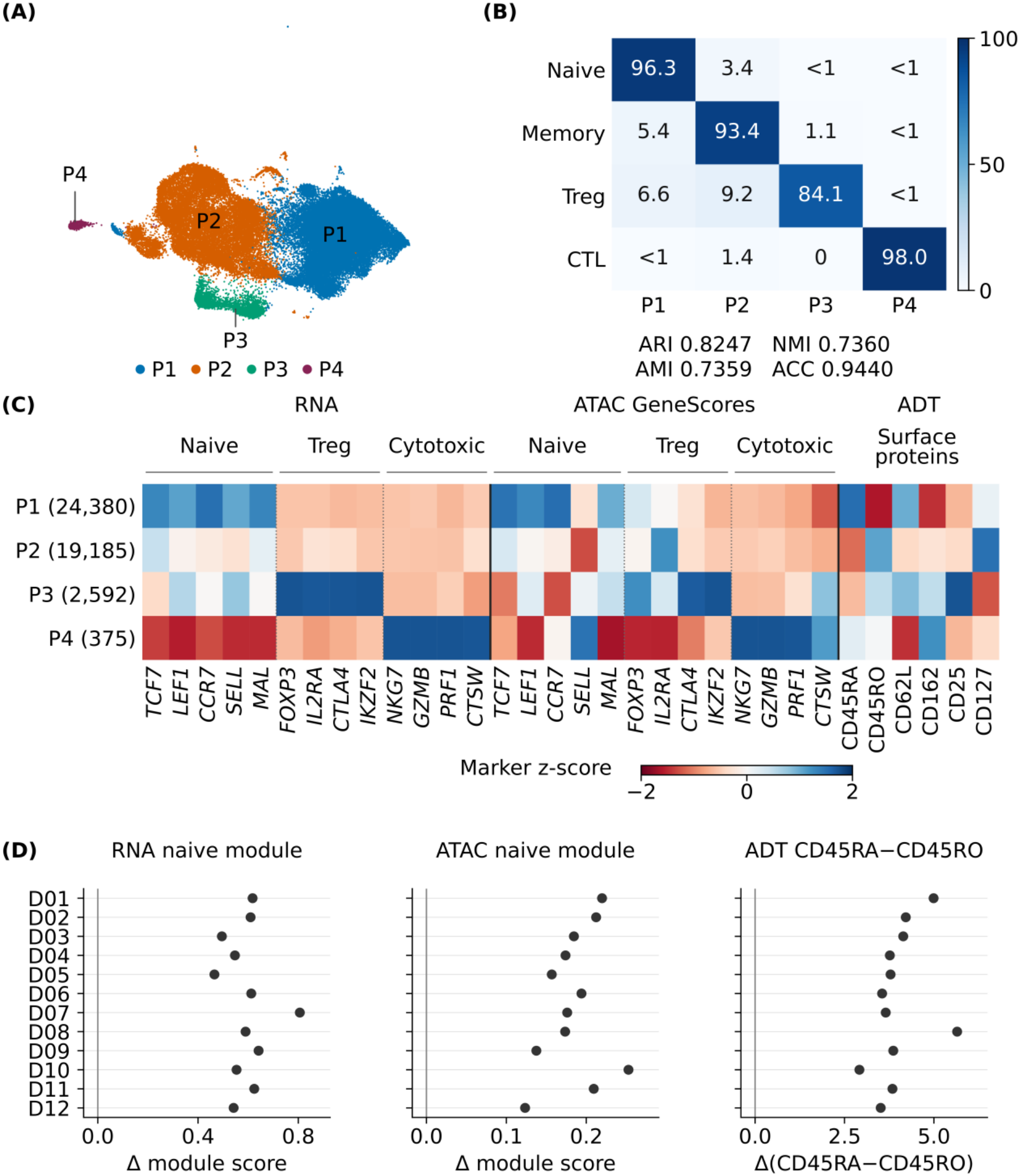
Tri-modal characterization of CD4-related cell communities. (A) UMAP of the selected GatorAnchor representation for 46,532 ALTRA TEA-seq cells, colored by predicted communities P1–P4. (B) Correspondence between predicted communities and the four reference categories. Values are row-normalized percentages; ARI, NMI, AMI, and ACC summarize overall agreement. (C) Donor-balanced molecular profiles of RNA markers, ATAC gene scores, and ADT features across the four predicted communities. (D) Within-donor differences between predicted P1 and P2 for the RNA naive module, ATAC naive module, and the ADT CD45RA–CD45RO score. Each point represents one donor (*n* = 12); positive values indicate higher scores in P1 than in P2.

Observed molecular profiles provided support for these community characteristics (**Figure 9C**). To balance donor contributions, measurements were first averaged within each donor–community group and then equally across the 12 donors. P1 showed higher naive-associated RNA expression of *TCF7, LEF1, CCR7, SELL*, and *MAL* than P2, together with higher CD45RA and CD62L ADT signals, whereas P2 showed higher CD45RO. ATAC gene scores for *TCF7, LEF1, CCR7*, and *MAL* were also highest in P1. P3 had the highest RNA means for *FOXP3, IL2RA, CTLA4*, and *IKZF2*, together with the highest CD25 and lowest CD127 ADT signals, consistent with a Treg-associated molecular profile. P4 showed the highest RNA and ATAC gene-score means for *NKG7, GZMB, PRF1*, and *CTSW*, supporting cytotoxic-associated features. These patterns retained modality-specific differences; for example, ATAC *IL2RA* was higher in P2 than in P3, despite the higher RNA *IL2RA* signal in P3.

We examined whether the molecular distinction between P1 and P2 was consistently observed across individual donors (**Figure 9D**). For each donor, we compared the mean molecular scores of cells assigned to P1 and P2. Across all 12 donors, P1 showed higher RNA and ATAC naive-associated module scores and a higher ADT CD45RA–CD45RO score than P2. These consistent within-donor differences support the molecular distinction between P1 and P2 across transcriptional, chromatin-accessibility, and surface-protein measurements.

## Methods

### Modality-specific representations and initial observation graph

Consider *n* cells jointly profiled for RNA and a second molecular modality. Let

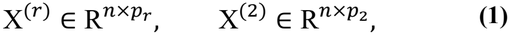

denote the RNA and second-modality data matrices, where the second modality denotes either antibody-derived tag (ADT) abundance or chromatin accessibility measured by ATAC sequencing (ATAC-seq).

Accordingly, GatorAnchor learns a *d*_*z*_-dimensional integrated representation for each cell. The learned cell representations are collected in the matrix:

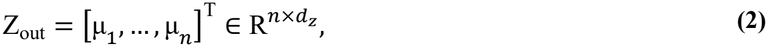

where μ_*i*_ is the fused latent mean for cell *i*. During training, stochastic latent sample z_*i*_ is used for reconstruction, whereas deterministic mean μ_*i*_ is used for edge discrimination, graph refinement, and downstream analyses.

For fully paired data, let M = {*r*, 2} denote the modality index set, where *r* denotes RNA and 2 denotes the second modality. The two modalities are processed independently. Measurements are normalized and log-transformed, after which the retained features are standardized and separately projected using modality-specific principal component analysis (PCA). The generated PCA score matrices can be written as:

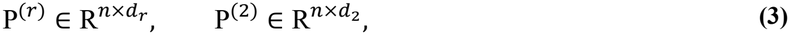

where *d*_*r*_ and *d*_2_ are the retained PCA dimensions for RNA and the second modality, respectively. These scores provide the features for graph construction and the reconstruction targets for the modality-specific decoders. Reconstruction therefore operates in PCA space.

To construct the initial observation graph, the two modality-specific PCA score matrices are directly concatenated:

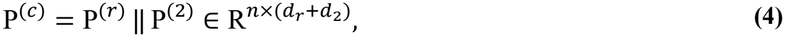

where ‖ denotes feature-wise concatenation. Note that no additional block-wise normalization or modality reweighting is applied at this step. The resulting geometry therefore depends on the magnitudes of the modality-specific PCA scores.

For cells with nonzero concatenated PCA score vectors, cosine distance is defined as:

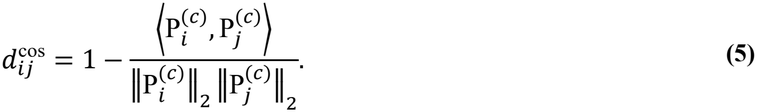

For each cell *i*, let 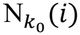 sdenote its *k*_0_ nearest non-self neighbors under this distance, where 1 ≤ *k*_0_ ≤ *n* − 1. The directed edge weights are defined as:

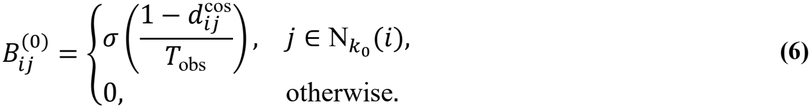

Here, *σ*(*x*) = (1 + *e*^−*x*^)^−1^ is the logistic sigmoid function, and *T*_obs_ > 0 controls the mapping from cosine similarity to edge weight. Neighbor selection depends only on cosine distance and is independent of *T*_obs_. For fixed nonzero similarities, decreasing *T*_obs_ moves positive-similarity weights toward 1 and negative-similarity weights toward 0. Increasing *T*_obs_ moves the weights of selected edges toward 1/2, whereas unselected relations retain zero weight. These values are used as similarity-based propagation weights rather than calibrated edge probabilities.

The directed graph is symmetrized using:

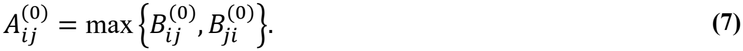

This operation forms the union of the directed neighborhoods, retaining an undirected edge whenever either cell selects the other. Because cosine similarity is symmetric, reciprocal selected relations have identical weights. The number of neighbors after symmetrization may exceed *k*_0_.

The symmetrically normalized propagation matrix is then obtained as:

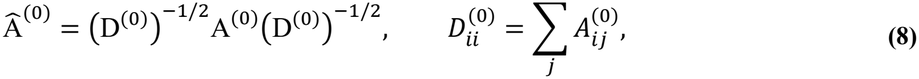

where D^(0)^ is the diagonal weighted-degree matrix. Self-neighbors are excluded during graph construction, and no self-loops are added during normalization. Thus, 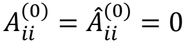.

The normalized graph is used to incorporate neighborhood information into each modality-specific representation. For each *m ∈* M, the encoder input under propagation graph Â^(*g*)^ is defined as:

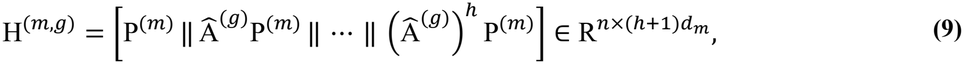

where *d*_*m*_ is the PCA dimension of modality *m*, and *h* ≥ 0 is an integer specifying the maximum propagation order. The zero-order block retains each cell’s own unpropagated PCA scores. For *t* = 1, …, *h*, the block 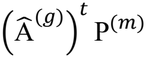 represents *t* successive steps of degree-normalized graph propagation. Both modalities use the same propagation matrix while retaining their separate feature blocks.

Here, *g* = 0 denotes the initial graph used in Stage A, whereas *g* =*\** denotes the refined graph constructed after warm-up. After refinement, the modality-specific inputs H^(*m,\**)^ are recomputed using Â^*\**^, which remains fixed throughout Stages B and C. The resulting graph-augmented feature matrices are supplied to the shallow branches of the modality-specific probabilistic encoders.

### Probabilistic graph encoding and disagreement-adjusted multimodal fusion

Given the graph-augmented features defined above, GatorAnchor learns modality-specific Gaussian representations in a shared *d*_*z*_-dimensional latent space. For each modality *m ∈* M, two linear heads map the graph-augmented features to shallow mean and log-variance representations:

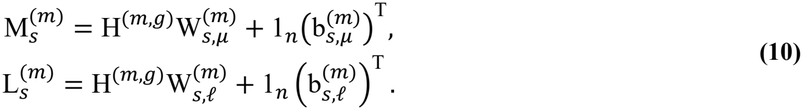

Both output matrices belong to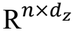. The trainable weight matrices belong to 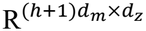, the bias vectors belong to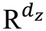, and 1_*n*_ is the length-*n* vector of ones.

During Stage A, only the shallow branch is active, such that the modality-specific Gaussian-parameter outputs are 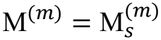 and 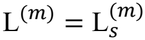, computed using the initial graph. After graph refinement, the shallow inputs are recomputed using Â^*\**^. During Stages B and C, a residual graph convolutional network (GCN)^28^ branch additionally processes the modality-specific PCA representation P^(*m*)^ on this fixed refined graph.

Let 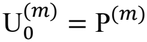, and let 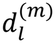 denote the width of GCN layer *l*, with 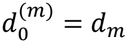. For *l* = 1, …, *L*, the residual branch computes:

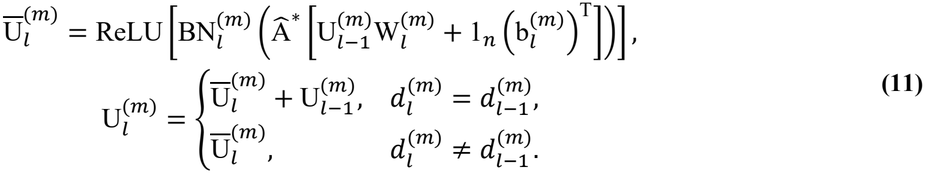

Here, *L*_*G*_ is the number of GCN layers, BN denotes batch normalization, 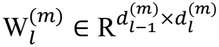, and 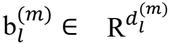. An identity residual connection is applied only when the input and output dimensions match. The branch uses the refined propagation matrix without adding self-loops.

Two linear heads map the final representation to residual corrections of the Gaussian parameters:

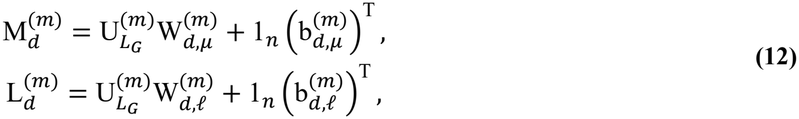

where 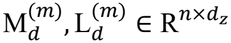.

During Stages B and C, the shallow representations and residual corrections are combined as:

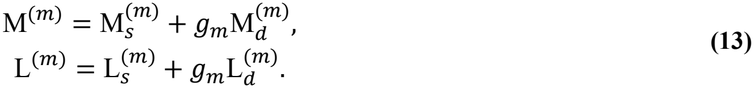

The modality-specific scalar *g*_*m*_ *∈* R is a trainable residual scale shared by the mean and log-variance corrections. It is initialized to zero and remains unconstrained during optimization. The corresponding variance is obtained by exponentiating the combined log variance.

Let 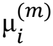 and 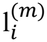 denote the transposed *i*th rows of M^(*m*)^ and L^(*m*)^, respectively. The modality-specific

Gaussian expert for cell *i* can be written as:

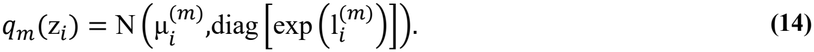

Dependence on the observed measurements, propagation graph, and learned model parameters is suppressed in this notation.

The two modality-specific Gaussian experts are combined using a product-of-experts (PoE) rule^29,30^, with a standard normal prior as an additional expert:

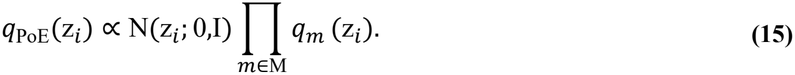

The coordinate-wise fused precision, mean, and variance can be written as:

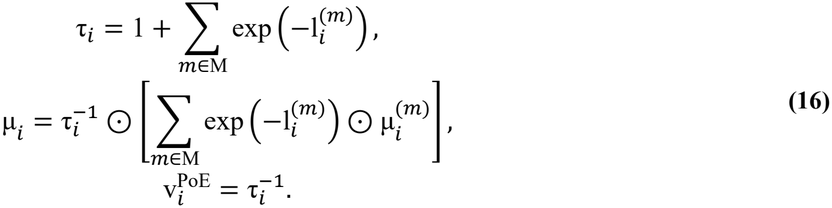

Here, 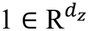, ⊙ denotes element-wise multiplication, and exponentiation and reciprocals are evaluated element-wise. Lower learned variance gives an expert greater relative influence within the corresponding latent coordinate. The prior contributes unit precision and a zero-mean reference to the fusion.

Because the modality-specific encoders share a graph-derived neighborhood structure, PoE is used operationally as a precision-weighted fusion rule. The fused distribution is not interpreted as a posterior derived under conditional independence of the modalities.

The PoE variance depends on the expert variances and does not directly reflect differences between their means. Thus, two low-variance experts can yield a small PoE variance despite substantial mean disagreement. GatorAnchor quantifies this disagreement using the coordinate-wise dispersion of the modality-specific means:

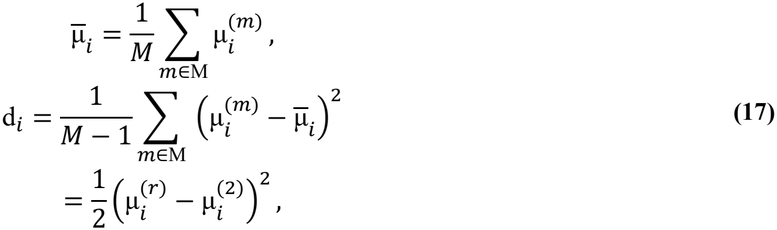

where squares are element-wise and the final equality follows from *M* = 2. This quantity is a deterministic disagreement measure computed from the two learned means, expressed in sample-variance form.

The fused variance is adjusted as:

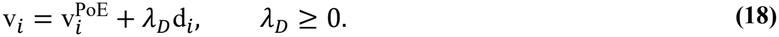

The adjustment adds a nonnegative contribution to each variance coordinate. When the modality-specific means coincide, d_*i*_ = 0 and 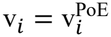 exactly. For fixed encoder outputs, this adjustment leaves the fused mean unchanged. The resulting distribution can be written as:

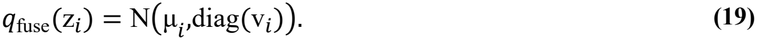

The additional term is a model-defined adjustment to latent dispersion; the adjusted covariance is not interpreted as a calibrated posterior covariance.

During training, reconstruction uses a reparameterized sample:

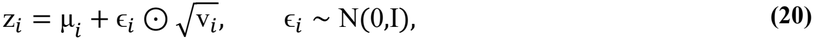

where the square root is element-wise. The deterministic fused mean μ_*i*_ is used for edge discrimination, graph refinement, and downstream analyses. Although the variance adjustment does not directly alter this mean, it can affect the learned representation through gradients from the reconstruction objective.

The Kullback–Leibler regularization defined below is applied separately to the modality-specific distributions *q*_*m*_, rather than to the disagreement-adjusted fused distribution *q*_fuse_.

### Self-supervised reconstruction and neighborhood discrimination

GatorAnchor learns the probabilistic representations through PCA-space reconstruction, neighborhood discrimination, and modality-specific Kullback–Leibler (KL) regularization. Throughout this subsection, M = {*r*, 2} and *M* = |M| = 2. Reconstruction uses reparameterized samples from the fused latent distributions, whereas neighborhood discrimination uses deterministic latent means.

Let

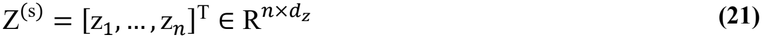

denote the matrix of fused latent samples defined above. For each modality *m ∈* M, a linear decoder reconstructs the corresponding PCA scores:

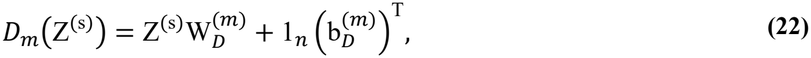

where 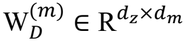 and 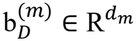 are trainable parameters, and 1*n* is the *n*-dimensional vector of ones.

The sampled reconstruction loss is:

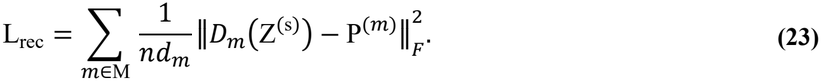

This expression is a Monte Carlo estimate of the expected reconstruction loss under the fused latent distributions. Each modality contributes its mean squared error per cell and retained PCA coordinate. The reconstruction targets are the modality-specific PCA scores.

A shared edge decoder learns to discriminate the directed neighborhood relations used to construct the initial observation graph. For a given representation, let 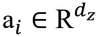 denote either the fused mean μ_*i*_ or a modality-specific mean 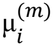. Both cells in a pair are represented using the same choice of latent representation. For an ordered cell pair (*i, j*), the pairwise feature vector can be written as:

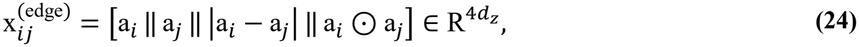

where ‖ denotes concatenation, |⋅| denotes element-wise absolute value, and ⊙ denotes element-wise multiplication. The decoder computes:

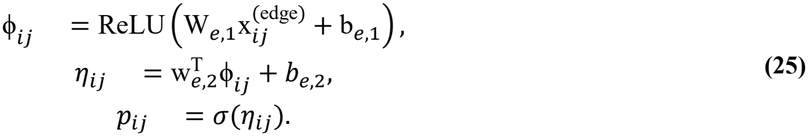

For hidden width 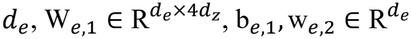, and *b*_*e*,2_ *∈* R. Here, ϕ_*ij*_ is the hidden pair representation, *η*_*ij*_ is the logit, and *σ* is the logistic sigmoid function. All decoder parameters are shared across the fused and modality-specific representations. Because a_*i*_ and a_*j*_ occupy distinct positions in the input, the decoder is not constrained to satisfy *p*_*ij*_ = *p*_*ji*_. The positive-pair set can be written as:

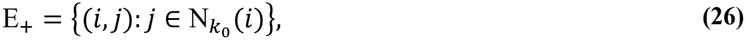

where 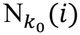 is the directed neighborhood defined before symmetrization of the initial graph. For each positive pair (*i, j*), the source cell *i* is retained and a destination 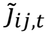 is sampled uniformly from {1, …, *n*} to form a pair assigned label 0. The labeled multiset used at epoch *t* is:

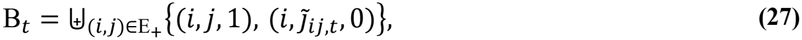

where ⨄ denotes multiset union, so repeated observations are retained. Self-pairs and existing positive relations are not excluded from negative sampling. Consequently, the same ordered pair can occur with both labels. The labels identify observed neighborhood pairs and uniformly sampled destination pairs; a label of 0 does not establish biological dissimilarity. Each positive observation contributes one negative observation, giving |B_*t*_| = 2|E_+_| = 2*nk*_0_.

For a fixed choice of latent representation, the edge-discrimination loss can be written as:

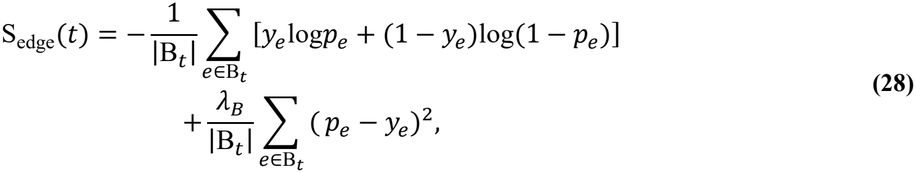

where *e* indexes a labeled observation, *y*_*e*_ *∈* {0,1} is its assigned label, *p*_*e*_ is the decoder score for its ordered cell pair, and *λ*_*B*_ ≥ 0 controls the binary Brier term^31^. The first term is binary cross-entropy, and the second penalizes squared deviations between the scores and assigned labels. These losses are evaluated under the constructed sampling distribution. The score *p*_*ij*_ is therefore used as a measure of edge support for this discrimination task; the training objective alone does not establish calibration to biological cell–cell connectivity.

The same labeled multiset B_*t*_ is used for the fused and modality-specific representations. Their losses are combined as:

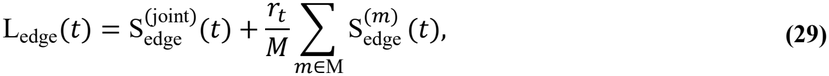

where the joint term uses μ and the modality-specific terms use μ^(*m*)^. The nonnegative coefficient *r*_*t*_ increases from zero to its prescribed terminal value during Stage A and remains at that value throughout Stages B and C. The positive-pair set E_+_ remains defined by the original directed neighborhoods after graph refinement. Thus, refinement changes the propagation graph while preserving the source of positive supervision for the discrimination task.

Each modality-specific Gaussian distribution is regularized toward the standard normal distribution:

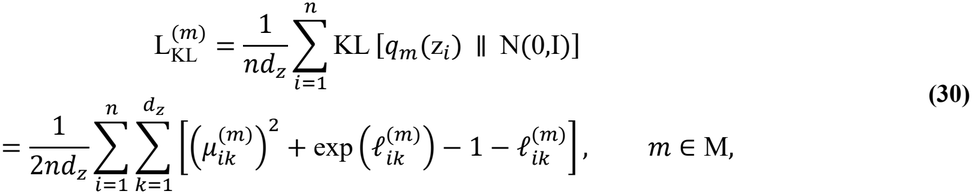

where 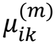 and 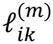 denote the *k*th coordinates of the modality-specific mean and log-variance vectors.

This term is averaged over cells and latent coordinates. The complete training objective is:

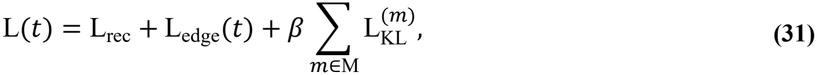

where *β* ≥ 0 controls modality-specific Gaussian regularization. The reconstruction and combined edge losses have unit coefficients, and the modality-specific KL terms are summed. This is a composite objective for stochastic representation learning. KL regularization is applied to *q*_*m*_ rather than to the fused distribution *q*_fuse_.

At the end of Stage A, the fused means are computed in evaluation mode. Together with the fitted edge decoder, they provide the decoder-derived evidence for the one-time graph-refinement procedure described below. The refined graph remains fixed throughout Stages B and C, while the edge decoder continues to be optimized through L_edge_(*t*). To complement the support scores, the following subsection defines a local score-variance proxy based on perturbations of the fitted final-layer weights.

### Post-hoc last-layer score-variance approximation

At the end of Stage A, the edge decoder is evaluated using the fused latent means, with the encoders, hidden features, and output bias fixed. Let 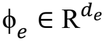 denote the hidden representation of an ordered pair *e* = (*i, j*) and ŵ the fitted final-layer weights. The reference multiset R_*L*_ contains the ordered candidates defined below and one additional pair (*i, J*_*e*_) for each candidate, where *J*_*e*_ is sampled uniformly from all *n* cells. Repeated pairs, self-pairs, and candidate-pair collisions are retained. The stabilized BCE-based curvature matrix can be written as:

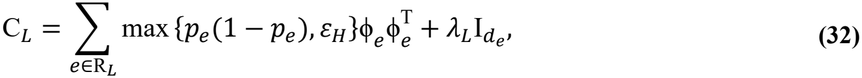

where *ε*_*H*_, *λ*_*L*_ > 0. This summed surrogate excludes the Brier term and is distinct from the Hessian of the complete training objective.

Under the auxiliary local perturbation model 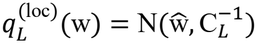, the first-order delta approximation motivates:

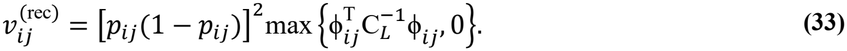

The nonnegative clipping handles numerical error. This reference-dependent proxy measures local score sensitivity and can be small for saturated outputs; it is not a calibrated posterior predictive variance or evidence that an edge is correct.

### Uncertainty-weighted evidence fusion and one-time graph refinement

For fully paired data, directed candidate proposals are concatenated in the following order:

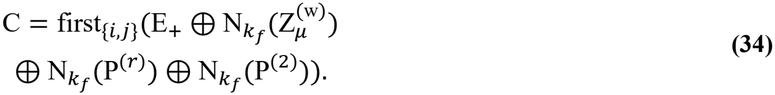

Here, E_+_ is the original directed neighborhood list, 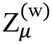 contains the evaluation-mode fused means at the end of Stage A, and ⊕ denotes list concatenation. The operator 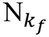 requests *k*_*f*_ + 1 cosine nearest neighbors per cell and discards the first returned neighbor. The operator first_{*i,j*}_ retains the first proposal for each unordered pair, preserving that proposal’s direction; the retained candidates are then sorted by unordered-pair key. Only this retained direction is scored by the decoder, and each unordered pair contributes once to candidate statistics. Original neighborhood pairs undergo the same fusion and pruning as new proposals.

For observed modalities *m ∈* M, define

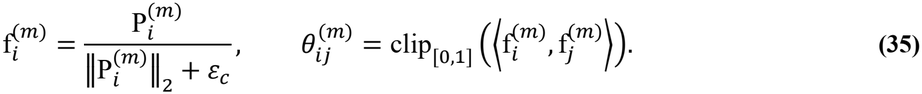

The branch precision summaries can be written as:

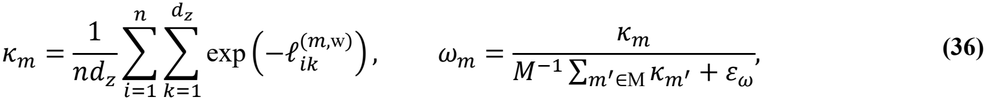

where 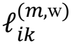 is the evaluation-mode log variance at the end of Stage A and *M* = |M| = 2. For sources S = {*r*, 2,rec}, the raw penalties and decoder support are:

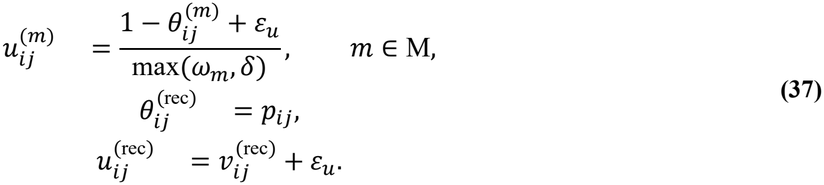

Each source is normalized over the common candidate pool:

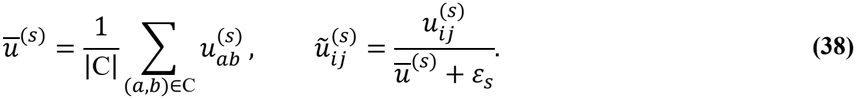

This normalization makes penalties relative within each source. The source-wise factor max(*ω*_*m*_, *δ*) cancels apart from its interaction with *ε*_*s*_. The fusion coefficients are

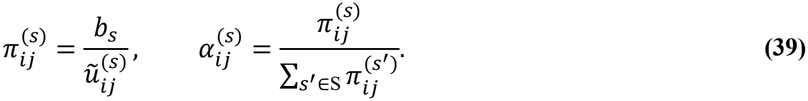

The fixed positive baseline coefficients satisfy *b*_*r*_ = *b*_2_ = *b*_obs_, with a separate *b*_rec_; these coefficients are independent of *ω*_*m*_.

The fused support, weighted disagreement, and composite penalty are:

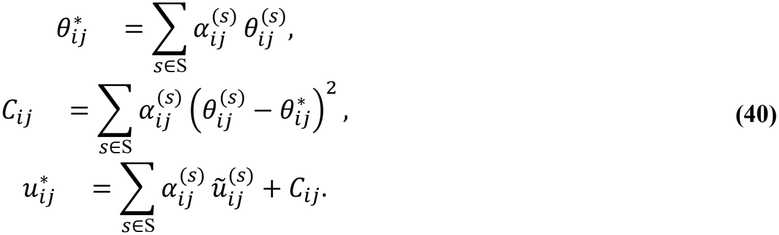

The conflict term measures disagreement under the fusion weights and can be small when one source dominates. These penalties and weights are algorithmic quantities, not posterior precisions or variances from a unified probabilistic model.

To characterize edge disagreement in the dual-modality analyses, define observation-only weights and consensus as:

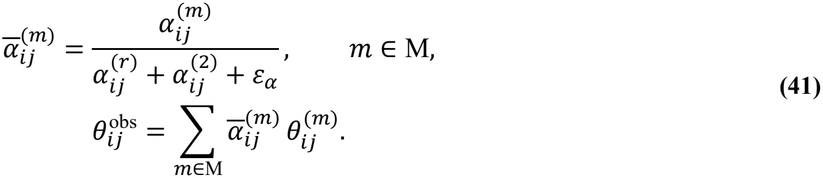

Modality conflict (also termed observation conflict) and reconstruction conflict are defined as:

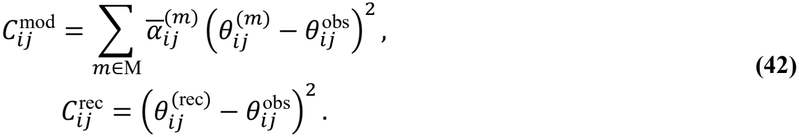

Here, reconstruction conflict quantifies decoder–observation disagreement in edge support, rather than molecular reconstruction error. These diagnostics are distinct from latent-mean disagreement d_*i*_ and total cross-source conflict *C*_*ij*_. Cell-level conflict scores and source weights are unweighted arithmetic means of the corresponding edge-level values over all incident candidates in C before pruning.

Let 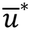 be the mean of 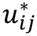 over C, and 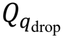 its empirical quantile, computed by linear interpolation. For *T*_unc_ > 0, the retained weights are:

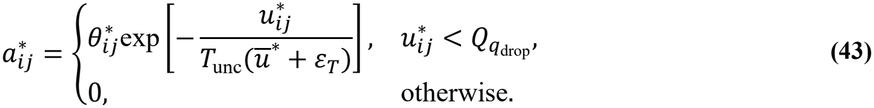

Here, *q*_drop_ is the quantile level, not the fraction removed; candidates tied at the threshold are discarded. Set 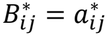 for (*i, j*) *∈* C and zero otherwise. The refined propagation matrix can be written as:

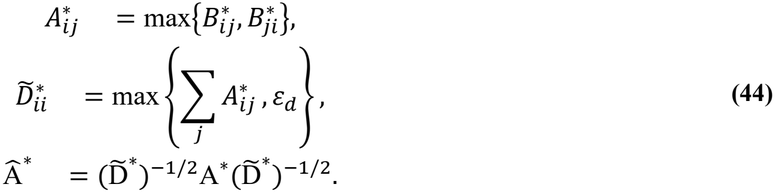

Thus, symmetrization follows directional scoring, fusion, and pruning. The degree floor leaves zero-degree rows zero; no self-loops or rescue edges are added. The inputs H^(*m,\**)^ are recomputed using Â^*\**^, which remains fixed throughout Stages B and C.

All *ε* constants in this subsection and *δ* are positive numerical stabilizers.

### Three-stage optimization curriculum

GatorAnchor minimizes the composite objective L(*t*) in three sequential stages, with one full-batch update per epoch and a single graph-refinement step between Stages A and B.

In Stage A, the residual GCN branch is bypassed. The shallow mean and log-variance heads, modality-specific reconstruction decoders, and shared edge decoder are trained on Â^(0)^ for *E*_pre_ epochs. The modality-specific edge-loss coefficient can be written as:

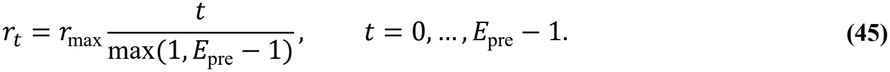

Here, *r*_max_ is the target weight of the average modality-specific edge-discrimination loss. At the end of Stage A, evaluation-mode fused means and modality-specific log variances, together with the fitted edge decoder, are used to construct Â^*\**^ once. The multi-hop inputs H^(*m,\**)^ are then recomputed on this refined graph.

Stage B activates the residual GCN branch and freezes only the parameters of the shallow mean and log-variance heads. The residual branch, including its output heads and gates, the reconstruction decoders, and the edge decoder remain trainable. The shallow heads receive the recomputed inputs, with dropout active during training. Stage C unfreezes the shallow heads and jointly fine-tunes all model parameters on the same refined graph.

The post-refinement budget *E*_ref_ is allocated as:

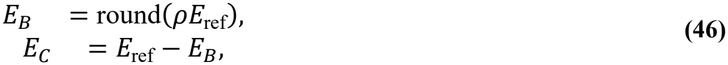

where *ρ ∈* [0,1] controls the allocation to Stage B. Throughout Stages B and C, *r*_*t*_ = *r*_max_ and Â^*\**^ remains fixed. Positive pairs for neighborhood discrimination remain the original directed neighborhood relations E_+_ throughout all three stages.

Each stage uses a newly initialized Adam optimizer, resetting optimizer states at stage transitions. Stages A and B use learning rate *η*_0_, and Stage C uses *c*_ft_*η*_0_, where *c*_ft_ is the fine-tuning multiplier. The final fused latent means are obtained in evaluation mode for downstream analyses.

### Application of GatorAnchor to incomplete and mosaic modalities

To apply GatorAnchor to incomplete and mosaic molecular measurements, we used a mask-aware implementation in which modality availability was tracked separately for each cell. Feature selection, scaling and PCA were fitted separately using only cells with the corresponding modality observed; missing PCA rows were zero-filled before propagation and tracked by presence masks. The PoE fusion included the prior once and only the experts corresponding to observed modalities. Unlike the sample-variance formulation above, mean disagreement was computed using the coordinate-wise population variance of the observed expert means, giving zero for cells with only one observed modality. Reconstruction and KL losses were averaged over observed cells and output dimensions within each modality and then summed. Modality-specific edge tasks required both positive-edge endpoints to have the corresponding modality observed, with negative endpoints sampled from cells in which that modality was observed; the fused task used all cells.

The initial graph united modality-specific observed-cell neighborhoods and scored edges using modalities shared by their endpoints. Refinement added fused-latent neighbors and renormalized weights over the available evidence sources, allowing decoder-only evidence for pairs sharing no observed modality.

Masks were shared across methods and retained at least one modality per cell. Reciprocal masks divided unpaired cells approximately equally between modalities; type-block masks targeted the five smallest classes that contained at least max(20, ⌊0.002*n*⌋) cells per selected dataset. Clustering used K-means with 50 initializations, with the number of clusters set to the number of reference classes. Reference labels specified the masks, cluster counts and evaluations but were not used as inputs to representation learning. Mask-aware PCA concatenated zero-filled modality PCs and presence indicators before applying PCA to at most 50 dimensions; the observed-RNA baseline used RNA PCs and its presence indicator.

Normalized robustness was rAUC = trapz(ARI(*r*), *r*)/[*r*_max_ARI(0)], where *r* denotes the hidden-cell fraction and ARI was averaged over masks and seeds within each dataset. Bidirectional nearest-centroid transfer used latent coordinates standardized over eligible unpaired cells and classes present in both modality groups; chance-adjusted balanced accuracy was (BA − 1/*C*)/(1 − 1/*C*) for *C* shared classes. Type-block F1 used global Hungarian matching, treating the hidden class as positive and counting errors across all cells.

Neighborhood purity measured the same-class fraction among 15 nearest neighbors, excluding self. Reference transfer used seed-matched full-data embeddings, with centering, global-RMS scaling and orthogonal Procrustes alignment fitted on non-hidden cells, followed by 15-neighbour majority voting with each query’s own counterpart excluded. Recovery inversely transformed decoder predictions through the fitted PCA and feature scaling, clipped negative values to zero, and correlated predicted and measured hidden-cell mean log-normalized profiles across retained second-modality features. Type-block metrics were averaged over seeds and then equally over blocks within each assay.

### Application of GatorAnchor to three molecular modalities

To apply GatorAnchor to fully paired RNA, ATAC, and ADT measurements, we adapted the dual-modality formulation to a three-modality setting. In this subsection, M_3_ = {*r, a, p*} and *M* = 3, where *r, a*, and *p* denote RNA, ATAC, and ADT, respectively. Each modality was processed separately to obtain PCA scores 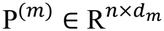 and assigned a modality-specific probabilistic encoder and linear reconstruction decoder. The initial observation graph was constructed from the concatenated representation P^(*r*)^‖P^(*a*)^‖P^(*p*)^, while all three modality branches shared the same propagation graph and edge decoder.

The PoE fusion rule was applied to the three modality-specific Gaussian experts, with the standard normal prior included once. Cross-modal mean disagreement can be computed as:

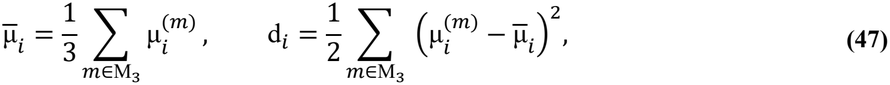

where squares are element-wise. The fused variance was adjusted as 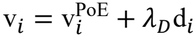, following the same formulation used in the dual-modality setting.

The reconstruction and KL sums were each multiplied by 2/*M*, preserving the total coefficients assigned to these modality-specific terms in the dual-modality objective. Defining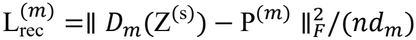, the training objective can be written as:

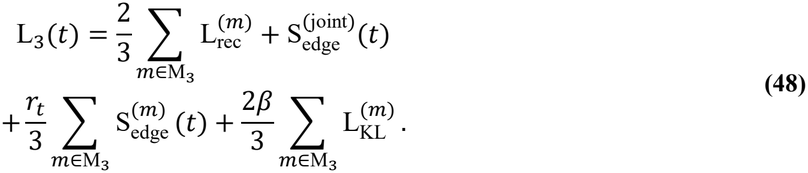

The individual reconstruction, edge-discrimination, and KL terms retained the definitions given above.

For graph refinement, candidate relations were assembled from the initial observation graph, fused latent means, and modality-specific PCA neighborhoods using the proposal and deduplication procedure described above. The evidence-source set was S_3_ = {*r, a, p*,rec}, comprising three observation-derived sources and the shared edge-decoder source. Source normalization, uncertainty-weighted fusion, conflict scoring, and edge filtering were then applied over this source set.

The observation-source baseline coefficients were set to *b*_*r*_ = *b*_*a*_ = *b*_*p*_ = 2*b*_obs_/*M*, preserving their sum at 2*b*_obs_. The decoder source used a separate coefficient *b*_rec_. The three-stage training schedule and one-time graph refinement were retained, with the final fused latent means used for subsequent analyses.

### Cell clustering and evaluation

Model-based clustering was performed on Z_out_ using mclust^32^, with the number of mixture components set to the number of reference classes for each dataset.

For datasets with reference annotations, clustering performance was evaluated using the adjusted Rand index (ARI), normalized mutual information (NMI), adjusted mutual information (AMI), and clustering accuracy (ACC). Let 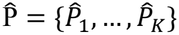 and R = {*R*_1_, …, *R*_*G*_} denote the predicted and reference partitions of the *n* cells, respectively. We define:

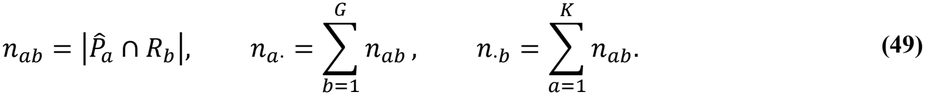

The ARI measures pairwise co-assignment agreement after correction for chance:

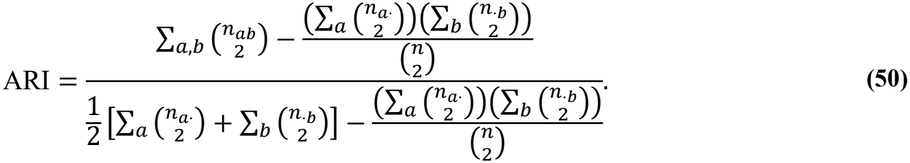

For the information-theoretic measures, the mutual information and partition entropies are given by:

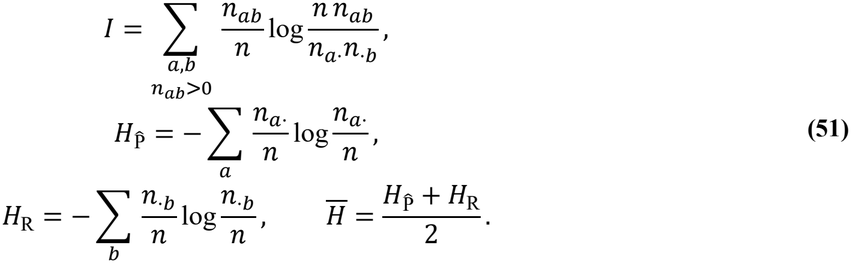

Using arithmetic-mean entropy normalization, NMI and AMI are given by:

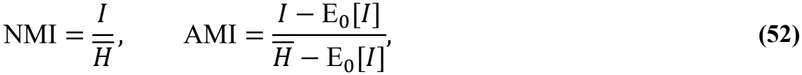

where E_0_[*I*] is the expected mutual information under the random-partition null model conditional on the observed cluster sizes.

Because cluster identifiers are arbitrary, ACC was computed after optimal matching between predicted clusters and reference classes:

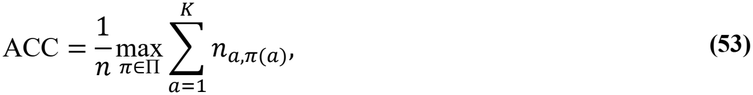

where Π is the set of admissible one-to-one label assignments between predicted clusters and reference classes. The optimal assignment was determined using the Hungarian algorithm.

### Implementation Details

GatorAnchor is implemented in Python 3.9.23 and PyTorch 2.4.1. Highly variable feature selection retains up to 4,000 RNA genes and 5,000 ATAC peaks; all ADT features are retained without feature selection. Up to 30 PCA components are used per modality, subject to the available sample and feature dimensions. The initial graph uses *k*_0_ = 15 and *T*_obs_ = 0.2, with propagation order *h* = 1. The latent dimension is *d*_*z*_ = 128, with three residual GCN layers of width 128 and edge-decoder hidden width *d*_*e*_ = 128.

Training uses full-batch Adam^33^ for 500, 150, and 100 epochs in Stages A, B, and C, respectively, with learning rates of 10^−3^, 10^−3^, and 10^−4^. Optimizer states are reset at stage transitions. The coefficients are *β* = 10^−5^, *λ*_*D*_ = 0.1, *λ*_*B*_ = 0.5, and *r*_max_ = 0.5.

Graph refinement is performed once after Stage A using *k*_*f*_ = 7 and *T*_unc_ = 1, retaining edges strictly below the 0.8 quantile of composite unreliability. The curvature damping and source coefficients are *λ*_*L*_ = 1, *b*_obs_ = 1.5, and *b*_rec_ = 0.4. The refined graph remains fixed thereafter. Final fused latent means are extracted in evaluation mode for downstream analyses.

For the three-modality application of GatorAnchor, the observation-source baseline coefficients were adjusted such that *b*_obs_ = 1.5 corresponded to *b*_*r*_ = *b*_*a*_ = *b*_*p*_ = 1.0, while *b*_rec_ = 0.4.

## Conclusion

We presented GatorAnchor, an uncertainty-aware framework for paired single-cell multi-omics integration that explicitly evaluates the reliability of cell–cell neighborhoods rather than treating graph structure as uniformly trustworthy. By combining modality-specific uncertainty with cross-modal and cross-source disagreement, GatorAnchor links probabilistic representation learning to refinement of the relationships used for graph propagation. Across ten RNA+ADT and RNA+ATAC datasets, GatorAnchor demonstrated strong performance in identifying cellular populations while providing interpretable diagnostics of edge reliability, modality conflict, and context-dependent evidence contribution. Biological analyses further showed that these diagnostics capture meaningful cross-modal measurement and neighborhood consistency. A tri-modal RNA–ATAC–ADT application further demonstrated that GatorAnchor can characterize CD4-related communities with molecular profiles and consistent within-donor molecular contrasts. Under incomplete and mosaic measurements, GatorAnchor preserved informative cell-population structure, while reliable cross-modal correspondence remained dependent on paired-cell anchors. These findings highlight neighborhood reliability as an important consideration in single-cell multi-omics integration and provide a framework for integrating heterogeneous molecular evidence while explicitly accounting for uncertainty and disagreement.

## Declarations

### Funding

Not applicable.

### Conflicts of interest/Competing interests

The authors declare that they have no competing interests.

### Ethics approval and consent to participate

Not applicable. This study involved secondary analysis of previously published and publicly available datasets. No new human participants were recruited, and no new biological samples or identifiable private information were collected by the authors as part of this study.

### Availability of data and material

All datasets analyzed in this study were previously published or publicly released; no new sequencing data were generated. The processed data used for the ten benchmark datasets listed in Table 1 are available from the NCBI Gene Expression Omnibus (GEO) under accession numbers GSE128639, GSE194122 (s3d6 RNA+ADT and s4d8 RNA+ATAC subsets), GSE158013 (RNA+ADT), GSE201402, GSE156478 (specifically the stimulated PBMC subseries GSE166188), and GSE126074 (adult mouse brain SNARE).

The 10Xmalt, 10X 1k PBMC, and PBMC unsorted 3k datasets are available from the corresponding 10x Genomics dataset pages. These datasets are distributed under the Creative Commons Attribution 4.0 International (CC BY 4.0) license.

The ALTRA TEA-seq dataset used for the three-modality analysis is publicly available through the Allen Institute Human Immune System Explorer (HISE; doi:10.57785/c0pq-s567), file altra_ARI-HC2_tea-seq.h5mu. The present analysis included 46,532 CD4-related cells from 12 donors.

### Code availability

The source code of GatorAnchor is available at https://github.com/zhangzh1328/GatorAnchor

### Consent for publication

Not applicable.

### Authors’ contributions

X.L., Y.L., and J.B. conceived the study and developed the methodology. Z.Z. and Y.L. implemented the software. X.L., Y.T., and Y.L. conducted the investigation, and Z.Z., Y.T., and Y.L. performed the formal analyses. Z.Z. curated the data, and Z.Z. and Y.L. prepared the visualizations. Z.Z., J.B., X.L., and Y.L. contributed to validation. J.B., X.L., and Y.L. supervised the study, with J.B. providing resources and project administration. Z.Z., X.L., and Y.L. drafted the manuscript. Y.T., J.B., X.L., and Y.L. reviewed and edited the manuscript. All authors read and approved the final manuscript.

### Clinical trial number

Clinical trial number: not applicable.

## Notes

### Competing Interest Statement

The authors have declared no competing interest.

